# Cellular basis of B_12_ uptake and remodelling in microalgae revealed using a novel bioassay

**DOI:** 10.64898/2026.08.07.738727

**Authors:** Ellen L. Harrison, Freddy Bunbury, Tobias Stadelmann, Andrew Sayer, Marcel Llavero-Pasquina, Konstantinos P. Papadopoulos, Katrin Geisler, Payam Mehrshahi, Matthew P. Davey, Alison G. Smith

**Author notes:** Author for Correspondence: Alison G. Smith.

## Abstract

- Vitamin B_12_, an essential micronutrient for many microalgae and humans, is synthesised only by certain prokaryotes. B_12_ is a complex tetrapyrrole that can exist in many forms (vitamers), some more bioavailable than others. Some microalgae are able to interconvert, or remodel, different B_12_ vitamers. As microalgae are important primary producers, it is crucial to understand how diverse microalgae acquire, utilise, and remodel this micronutrient.
- Through the development of a novel algal bioassay for B_12_ quantification that distinguishes between B_12_ vitamers with different lower axial ligands, and the generation of targeted knock-out lines, we characterised the role of proteins involved in algal B_12_ uptake and remodelling.
- We found that the previously characterised protein CoBalamin-Acquisition protein 1 (CBA1) is also necessary for the acquisition of pseudocobalamin, a less bioavailable form of B_12_. In addition, we provide the first experimental evidence that COBT is required for *Chlamydomonas reinhardtii* to remodel B_12_.
- We apply the algal B_12_ bioassay to show that the edible alga *Chlorella vulgaris* can accumulate pseudocobalamin but is unable to remodel it, highlighting the need for thorough investigation of the metabolic requirements and capabilities of microalgae, especially given the growing interest in microalgae-based food additives.

## Introduction

Vitamin B_12_ is a complex corrinoid molecule that acts as a cofactor for a range of enzymes, including cobalamin-dependent methionine synthase (METH). Its biosynthesis is confined to a subset of prokaryotes (Martens *et al*., 2002), and is not found in any eukaryote. Due to its costly synthesis, requiring over 20 catalytic steps, and its patchy distribution in the environment, it is perhaps surprising that so many microalgal species, characterised by their photosynthetic nature, are dependent on an exogenous supply of the vitamin (Croft *et al*., 2005; Tang *et al*., 2010; Helliwell *et al*., 2011; Lin *et al*., 2022). Indeed, vitamin B_12_ has been suggested as an ideal candidate to control microbial community interdependencies and nutrient exchanges (Sokolovskaya *et al*., 2020; Soto *et al*., 2023). Three B_12_-dependent enzymes are known in algal lineages, METH and methylmalonyl-CoA mutase (MCM), as in humans, along with type II ribonucleotide reductase (RNRII) (Helliwell, 2017). A study that combined literature search and laboratory culturing found that over 50% of the surveyed algae across all phyla (326 species) required an exogenous supply of B_12_ for growth in culture, and that this trait is scattered across algal phylogenetic lineages (Croft *et al*., 2005). Further work showed that B_12_ auxotrophy is due to the absence of a functional B_12_-independent form of methionine synthase (METE), the loss of which has occurred several times independently across the algal groups (Helliwell *et al*., 2011).

The relationship of microbes and vitamin B_12_ is further complicated by the fact that B_12_ is the general name for a range of cobalt-containing tetrapyrroles that can vary at the upper and lower axial ligand position (**Figure 1**). The upper-axial position is occupied by a methyl, hydroxy, deoxyadenosyl or cyano group (**Figure 1a**), and is involved in the cofactor’s radical chemistry. There is greater diversity at the lower axial ligand position, with three possible groups of compounds providing the base that coordinates to the cobalt atom: benzimidazoles, purines or phenols (**Figure 1b**). From the combination of various known upper and lower axial ligands there are thought to be more than 30 naturally occurring forms of B_12_ (Kennedy & Taga, 2020).

**Figure 1.**
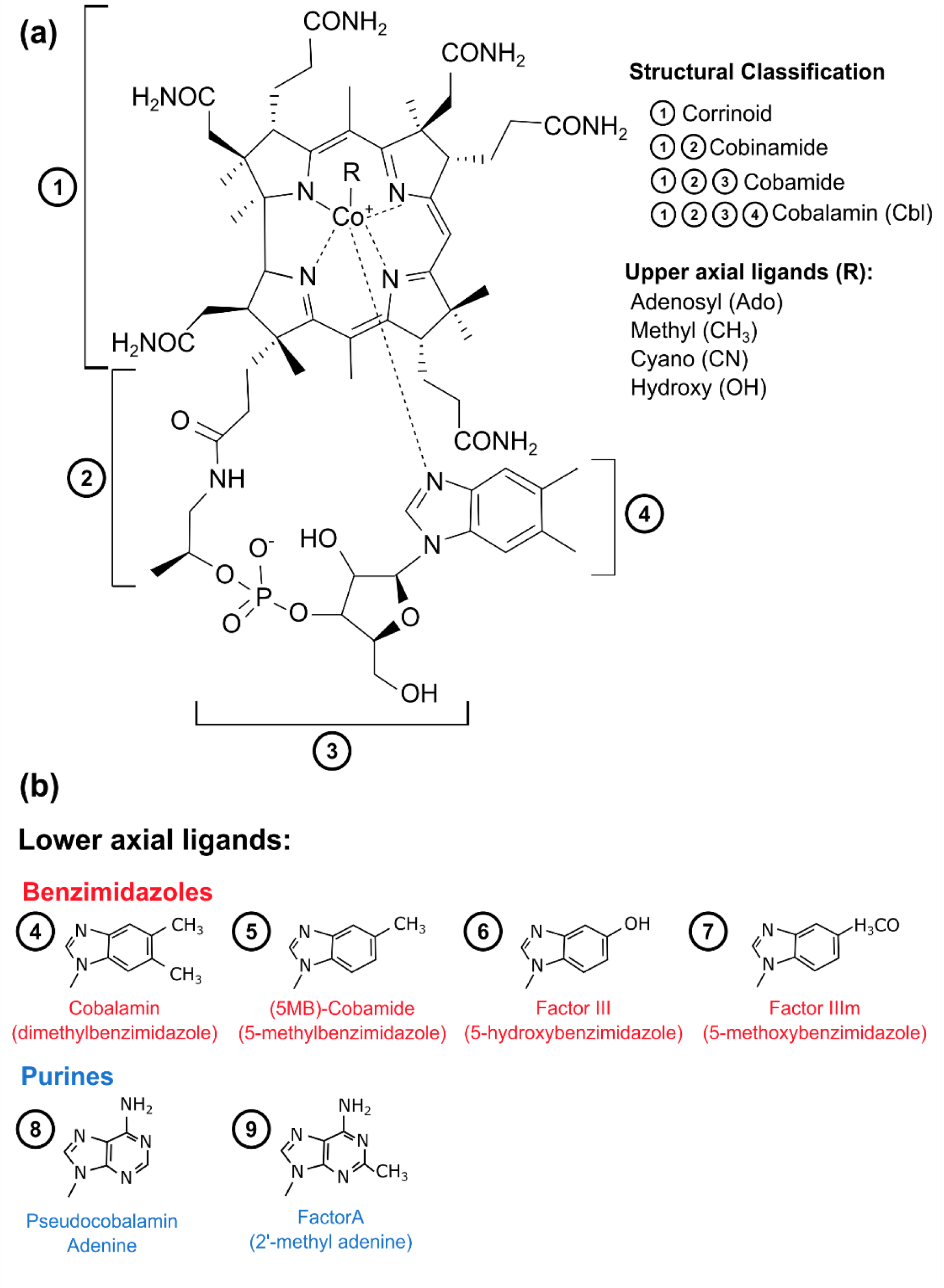
Structure of B_12_ vitamers. **a)** Structure of cobalamin, indicating the naming conventions of corrinoids more generally. (1) - corrinoid ring containing central cobalt ion; (2) – aminopropanol group; (3) – phospho-ribose; (4) base, lower axial ligand – in the case of cobalamin, dimethylbenzimidazole. Together (2), (3) and (4) constitute the nucleotide loop. R indicates the possible upper axial ligands to the central cobalt ion. **b)** Lower axial ligands can be benzimidazoles, purines or phenolic compounds (Shelton *et al*., 2018). The ones used in this study are depicted here.

In humans, the structure of the B_12_ vitamer can be central to its ability to act as a suitable cofactor for dependent enzymes and its recognition by B_12_-transport proteins (Stupperich & Nexø, 1991; Fedosov *et al*., 2007; Keller *et al*., 2018; Sokolovskaya *et al*., 2021). The lower axial ligand is particularly relevant for human nutrition, as those vitamers with a purine base, such as pseudocobalamin (**Figure 1b** structure 8), are less bioavailable than cobalamin, where dimethyl-5,6-benzamidazole (DMB) (**Figure 1b** structure 4) occupies the lower axial ligand position (Stupperich & Nexø, 1991; Sokolovskaya *et al*., 2021). In particular, intrinsic factor (IF), a protein that binds B_12_ in the intestine and facilitates its uptake by ileal cells, has a binding affinity six orders of magnitude weaker for pseudocobalamin and Factor A (**Figure 1b** structure 9), than for cobalamin (Fedosov *et al*., 2007).

In gram-negative bacteria, uptake of B_12_ is facilitated by components of the *btu* operon. *btuB* encodes a TonB-dependent protein needed to transport B_12_ across the outer membrane into the periplasm (Chimento *et al*., 2003). B_12_ is then bound by a high-affinity periplasmic binding protein, BtuF, which transfers it to the inner membrane ABC-transporter (BtuCD) (Schauer *et al*., 2008). In the human gut, some bacteria have been shown to use multiple transporters, giving them a competitive advantage to take up different B_12_ forms (Degnan *et al*., 2014). Much less is known about how microalgae accumulate B_12_ and in what forms. While the presence of pseudocobalamin has been detected within *T. pseudonana* cells (Heal *et al*., 2017), the mechanism of uptake was not investigated. Some of the earliest evidence of B_12_-binding proteins in microalgae was described by Pintner and Altmeyer (1979) who found that a range of microalgae released a protein into the medium that bound B_12_, and a protein that bound B_12_ with high affinity was isolated from the spent medium of the diatom *Thalassiosira pseudonana* (Sahni *et al*., 2001). More recent work has shown that a protein called <u>C</u>o<u>B</u>alamin <u>A</u>cquisition <u>P</u>rotein 1 (CBA1; Bertrand *et al*., 2012), is essential for B_12_ uptake in the diatom *Phaeodactylum tricornutum* and the green alga *Chlamydomonas reinhardtii* (Sayer et al., 2024). Through bioinformatic analysis, CBA1-like proteins were found in all major eukaryotic algal lineages, and in many cases its presence overlapped with organisms also having B_12_-dependent enzymes (Sayer *et al*., 2024).

As in humans, many microalgae also favour cobalamin and are unable to grow on pseudocobalamin (Helliwell *et al*., 2016). Several species of microalgae able to grow on pseudocobalamin if supplied with DMB (Helliwell *et al*., 2016), implicating remodelling of the cofactor to the bioactive cobalamin, in a way similar to many bacteria (Gray & Escalante-Semerena, 2009; Yi *et al*., 2012; Crofts *et al*., 2013; Shelton *et al*., 2018). In support of this, *C. reinhardtii,* which has remodelling capacity, encodes homologues of the bacterial CobT, which activates DMB, along with CobS and CobC involved in attachment of activated DMB to the cobamide precursor (Anderson *et al*., 2008). Since cyanobacteria synthesise pseudocobalamin rather than cobalamin, remodelling may be a way for microalgae to compete for this scarce micronutrient in the photic zone (Helliwell *et al*., 2016). The dissolved levels of B_12_ in aquatic environments can be highly variable, depending not only on the location but also the season (Ohwada & Taga, 1973; Panzeca *et al*., 2009; Sañudo-Wilhelmy *et al*., 2012; Bonnet *et al*., 2013). In the marine environment, reported levels of B_12_ were 0-87 pM in coastal regions compared to deeper oceanic values of 0-6.2 pM (Sañudo-Wilhelmy *et al*., 2014). In freshwater lakes, levels of B_12_ also varied with the season and depth with values from undetectable to 75 pM (Daisley, 1969; Cavari & Grossowicz, 1977). Most microalgae require a minimum of 10-100 pM to grow effectively in culture (Helliwell *et al.,* 2016), and B_12_ has been demonstrated to be limiting for phytoplankton growth in environmental amendment studies (Bertrand et al., 2007; Browning et al., 2017; 2018).

Here we investigate the physiology of algal B_12_ uptake and remodelling. First, we investigated the ability of two B_12_-dependent mutants from different algal lineages to grow on varying concentrations of B_12_ analogues, their ability to remodel the purinyl analogues, and the genes necessary for this process. We established a novel algal B_12_ bioassay that can distinguish between B_12_ vitamers that differ in their lower axial ligand. In parallel, we investigated both uptake and remodelling in the commercially relevant species *Chlorella vulgaris*. *C. vulgaris* is one of several algal species to have GRAS (generally recognised as safe) status, meaning it can be incorporated into human food (Wells *et al*., 2017; Neumann *et al*., 2018). Chlorella has been proposed as a future food (FAO. Fisheries, 2008; Parodi *et al*., 2018), based on its amino acid profiles, levels of omega-3 fatty acids, other micronutrients such as iron (Parodi *et al*., 2018) as well as a source of vitamin B_12_ (Watanabe *et al*., 2002). Chlorella products can contain highly variable amounts of B_12_. In one study of Chlorella supplements a range of <0.1 µg to 415 µg per 100 g dry weight was detected, with cobalamin found to be the predominant form (Bito *et al*., 2016). Since the B_12_ in algal cells is not produced by the alga itself, it is crucial to gain more information about the source of the vitamin in the supplements. We discuss the significance of our results both for microbial community dynamics, and for the use of microalgae in providing adequate nutrition, given the increasing the popularity of plant-based diets (Becker, 2007; Wells *et al*., 2017; Cole *et al*., 2018; Parodi *et al*., 2018).

## Materials and Methods

### Algal strain and media

The algal strains used in this work are detailed in **Table S1**, along with the media and growth conditions. All strains were maintained axenically and regularly checked for contamination by plating on LB, TY or marine broth media as appropriate. B_12_-dependent microalgal strains were supplemented with cyano-forms (**Figure 1a**) of the various B_12_ isoforms. The molecular mass varies slightly between vitamers, so where relevant either the molar concentration and/or the concentration in pg/ml is provided. Cyanocobalamin was obtained from Sigma-Aldrich (Dorset, UK). Other forms were obtained as described below.

### Algal culture monitoring

Algal growth was monitored using the optical density (OD) of the cultures at 730 nm on a CLARIOstar Plus plate-reader (BMG labtech). Cell counts for *C. reinhardtii* and *P. tricornutum* were performed using a Dual Threshold Coulter Counter Cell and Particle Counter (Beckman Coulter, Brea, USA) with a 70 µm probe, set to measure between 3-9 µm. Depending on the density of the cultures, between 10-100 µl of algal culture was added to 10 ml of fresh Isoton II diluent (Beckman Coulter, Brea, USA). Due to the small size of *C. vulgaris,* a haemocytometer and light microscope were used to count the cells.

### Algal growth on B_12_ analogues

The B_12_ analogues used in this work (**Figure 1b**) were a gift from colleagues Martin Warren (University of Kent, UK) and Bernhard Kräutler (University of Innsbruck, Austria). The experiment was conducted in 24-well plates with the conditions detailed in **Table S1** for each species, and the media supplemented with the stated B_12_ analogue concentration. The starting density of the *C. reinhardtii* cultures was adjusted to 3 x10^4^ cells/ml and for *P. tricornutum* to an OD_730_ of 0.01. *P. tricornutum* was also grown for one round of sub-culturing without B_12_ to deplete internal stores prior to the start of the experiment. Cultures were grown until stationary phase and the resulting OD_730_ normalised by dividing all values by the highest value reached for each species. In the case of *P. tricornutum* C9 at the lowest two concentrations of Factor A tested only two biological replicates were included due to measurement error, all others show results of biological triplicates.

### B_12_ uptake assay

B_12_ uptake was measured using the method detailed by Sayer *et al*. (2024) and was adapted for the inclusion of pseudocobalamin, and the use of the novel bioassay strain *C. reinhardtii* metE7. **Table S2** shows the density of the cells used for each strain of microalgae. The volume of cells required was added to 1.5 ml microcentrifuge tubes and centrifuged at 8000 rpm for 1 minute, then 12,000 rpm for 1.5 minutes to pellet the cells (Eppendorf^TM^ Benchtop Microcentrifuge 5424). The supernatant was removed and 1 ml of fresh media supplemented with B_12_ (concentrations in **Table S2**) was added to the tubes. If samples were incubated for more than 2 minutes, they were placed in the standard culturing conditions of the alga being tested, otherwise maintained at room temperature. Samples were inverted manually at the halfway point to aid in mixing. After the designated incubation period, samples were centrifuged as before. The supernatant was removed and retained for the assay as the media fraction. 1 ml water was added to the cell pellets and mixed. The cell and media fractions were then boiled for 25 minutes to release intracellular B_12_ into solution. Samples were either used in the bioassay straight away or frozen at-20 °C until use. The volume of sample used in the bioassay then depended on the concentration of B_12_ used in the initial incubation, typically it ranged between 100 and 500 µl. In the case of the *C. vulgaris* time course experiment, only two biological replicates were included at 10 minutes for the media fraction due to measurement error, all others show results of biological triplicates.

### *C. reinhardtii* metE7 B_12_ bioassay

*C. reinhardtii* metE7 was maintained as detailed in **Table S1**, then subcultured by transfer of 200 µl of stationary phase culture into 20 ml of TAP medium + 200 pg/ml cyanocobalamin and incubated for 4-5 days at 25°C, 110 rpm and a 16:8 light dark cycle (90 µE m^-2^s^-1^). The cell density was measured to calculate the necessary amount of culture required for the number of 24-well bioassay plates, with 2 ml per well to a final density of ∼60,000 cell/ml. The required volume of *C. reinhardtii* metE7 cells were spun in 1.5 ml Eppendorf tubes for 1 min at 8000 rpm, the supernatant was removed, and the pellet resuspended in fresh TAP (without B_12_) to remove extracellular B_12_, this process was repeated once more. The washed cells were added to the desired volume of fresh TAP and 1 ml aliquoted into each well, followed by addition of 1 ml of samples to be measured, along with a set of B_12_ standards. The plates were sealed and incubated at 25°C, 120 rpm, continuous light, 90 µE m^-2^s^-1^. After 4 days the cultures in the wells were mixed by pipetting and OD_730_ was measured as above. The resulting data was then analysed using four parameter logistic equation (Ritz *et al*., 2015), as detailed in supplementary materials.

### Generation of knock-out mutants by CRISPR-Cas9 gene editing

METE knockout lines were generated in *P. tricornutum* strain 1055/1 (**Table S1**) by CRISPR-Cas9 editing to introduce a nourseothricin cassette into the gene as previously described (Llavero-Pasquina et al., 2022). The sequence of the gRNAs are shown in **Table S3**. Transformants were cultured on selective media with 75 µg/ml zeocin, 300 µg/ml nourseothricin and 200 pg/ml cyanocobalamin and screened at the *PtMETE* locus using PCR with primers that differentially amplified the WT or mutated sequences (**Figure S1a & b**). Monoallelic lines were restreaked several times to obtain biallelic lines, which were then screened for cobalamin auxotrophy in liquid media without B_12_ supplementation (**Figure S1c**). Two lines, C9 and H9 were obtained, with C9 used for further experiments.

Gene-editing of the *COBT* gene in *C. reinhardtii* was carried out by introduction of a ribonucleoprotein complex, essentially as previously described (Ferenczi *et al*., 2021) with a few modifications, including the use of Cas9 nuclease rather than Cas12a. The target site in Exon 1 of the *COBT* gene (Cre12.g555500.t1.2) is shown in **Figure S2,** with the sequence of the single-stranded DNA template to introduce a stop codon in all possible reading frames. For RNP formation, 0.263 nmol Cas9 nuclease (New England Biolabs) was mixed with 0.789 nmol gRNA (**Table S3)** at 37 °C for 10 minutes. Cultures of *C. reinhardtii* metE4 were maintained in TAP + 200 pM cobalamin, 25 °C, continuous light (∼90 μmol m^-2^ s^-1^) until harvest at a density 1-5×10^6^ cells/ml, then cell concentration was adjusted to 2×10^6^ cells/ml in TAP + 200 pM cobalamin + 40 mM sucrose (TCS media). The RNPs were then mixed with 125 µl cells, and with 2.63 nmol single-stranded DNA template. Cells underwent electroporation with a BioRad GenePulser Xcell electroporator at 600 V, 50 µF, 200 Ω in a 4 mm cuvette. Immediately after electroporation, 800 µl TCS media was added to the reaction cuvette, followed by transfer to 5 ml of the same solution and incubation for 24 hrs at 120 rpm, 30 °C in low light conditions. The reactions were plated at different dilutions on TAP agar with 75 µg/ml kasugamycin, 50 µg/ml ampicillin and 1 nM cobalamin utilising a starch plating method, adapted from Ferenczi et al. (2021), where 800 µl of a 37.5% corn starch solution in TAP + 200 pM cobalamin was mixed with 200 µl of cells and added to the top of a TAP agar plate. The starch layer was air-dried in a sterile flow hood before sealing with parafilm. Plates were then incubated in continuous light, 25 °C until visible colonies could be picked. Individual colonies were picked into 200 µl TAP + 200 pM cobalamin in single wells of a 96-well plate and grown until stationary phase. The cells were then subcultured 1/200 in either TAP, TAP + 200 pM cobalamin or TAP + 200 pM pseudocobalamin + 1 µM DMB for screening (**Figure S3a**). A subset of lines that did not grow were genotyped by PCR to confirm the insertion of the stop codon (**Figure S3b**).

### Data analysis and visualisation

Data analysis and visualisation was carried out in Rstudio (version 4.4.3, https://www.R-project.org). The R package Tidyverse (Wickham *et al*., 2019) and ggpubr (Kassambara, 2025) was used to make the figures, along with the package drc (Ritz *et al*., 2015) for the 4-parameter logistic equation in the bioassay. The normality of the data was checked using the Shapiro test function in R, and then either the parametric or non-parametric version of a test applied as applicable. To perform a Dunn Test, the non-parametric post-hoc analysis of a Kruskal-Wallis test was carried out with the R package FSA (Ogle *et al*., 2022).

## Results

### Testing ability of model algae to utilise and remodel diverse B_12_ analogues

As shown previously by Helliwell *et al*. (2016), pseudocobalamin is orders of magnitude less bioavailable to a range of eukaryotic algae than cobalamin, including a B_12_-dependent *C. reinhardtii* metE7 strain, generated by experimental evolution (Helliwell *et al*., 2015). Other algae for which pseudocobalamin was found to be less bioavailable included the marine species *T. pseudonana*, *Ostreococcus tauri*, *Pavlova lutheri*, *Aureococcus anophagefferens,* and the freshwater species *Lobomonas rostrata*. Moreover, while some species were capable of restoring their growth on pseudocobalamin with the addition of DMB, such as *C. reinhardtii* metE7 and *P. lutheri*, some could not, including the diatom *T. pseudonana*. To investigate this further, we generated a B_12_-dependent *metE* mutant of the marine diatom *P. tricornutum* (*P. tricornutum* C9) by CRISPR-Cas9 as described in Methods. We used this, together with the *C. reinhardtii* metE7, as indicator strains to establish their preference for a range of B_12_ analogues. The strains were inoculated into fresh medium supplemented with these compounds at different concentrations. As well as pseudocobalamin, we tested Factor A, another purinyl lower axial ligand analogue, and three other benzimidazole-containing vitamers, along with cobalamin (**Figure 1b**). The growth of algal cultures was measured using OD_730_ after 4 days for *C. reinhardtii* metE7 and 15 days for *P. tricornutum* C9. The OD values were normalised by dividing all values by the highest optical density achieved by each organism.

*C. reinhardtii* metE7 showed a dose-dependent growth pattern on all the benzimidazole B_12_ analogues tested (**Figure 2a**). At the higher concentrations tested all benzimidazole analogues supported the growth to the same level. At the lowest concentration tested 0.05 nM, which is the most environmentally relevant, Factor III and Factor IIIm supported the growth the least well with a significant effect on the growth of *C. reinhardtii* metE7 by B_12_ form at 0.05 nM (Kruskal-Wallis test, *p*=0.02). As expected, there was negligible growth on pseudocobalamin unless supplied at the highest concentration (10 nM, ∼100x that found in the environment); this was also true for Factor A (**Figure 2a**). However, the addition of DMB to 1 nM of pseudocobalamin or Factor A was sufficient to restore the growth to the equivalent level of cobalamin (**Figure 2b**), due to remodelling.

**Figure 2.**
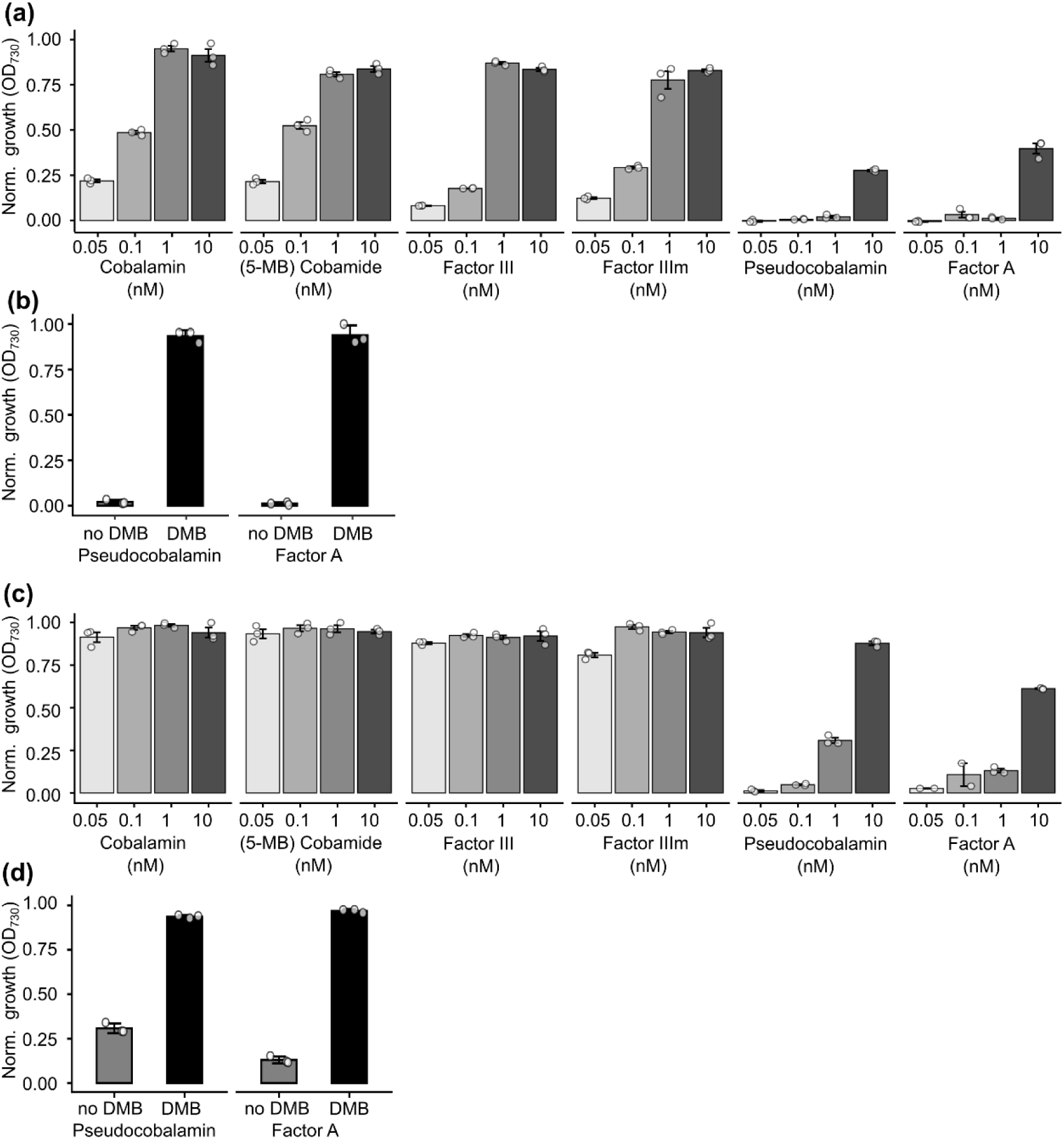
**Growth of B_12_-dependent microalgal mutants on various B_12_ analogues**. **a)** The B_12_-dependent *C. reinhardtii* metE7 mutant grown on a range of analogues at concentrations ranging from 0.5 nM to 10 nM. Cobalamin, Factor III and Factor IIIm have benzimidazole ligands, pseudocobalamin and Factor A have adenine ligands, and cobamide has no lower ligand. **b)** Growth of *C. reinhardtii* metE7 on 1 nM pseudocobalamin or Factor A with and without 1 µM DMB. B_12_ analogues. **c)** and **d)** shows the same as a) and b) respectively, but with the B_12_-dependent *P. tricornutum* strain C9. Species were grown until stationary phase (4 and 7 days, respectively), and normalised growth measured by taking the optical density at 730 nm (OD_730_) and dividing all values by the highest optical density achieved for each organism. n=3* ±SD. *See Materials and Methods.

Growth of the B_12_-dependent *P. tricornutum* C9 was supported by all the benzimidazole B_12_ analogues, and in contrast to *C. reinhardtii* metE7, there was substantial growth even at the lowest concentration used (**Figure 2c**). Although, at 0.05 nM B_12_ there was also a significant effect on the growth of *P. tricornutum* C9 depending on form B_12_ with benzimidazole lower axial ligands (Kruskal-Wallis test, *p*<0.05). Pseudocobalamin and Factor A did not support growth unless supplied at 10 nM (**Figure 2c**), suggesting that both can act as inefficient cofactors of METH if supplied in excess, with pseudocobalamin more effective than Factor A. Once again however, the inclusion of DMB to 1 nM pseudocobalamin or Factor A enabled growth, indicating that *P. tricornutum* C9 is also capable of remodelling (**Figure 2d**).

### Development of a new microbiological B_12_ bioassay

Historically several microbial strains have been used in bioassays to quantify B_12_ (Ford & Hutner, 1955; Hutner *et al*., 1956; Anderson, 1964; Kelleher & Broin, 1991; Raux *et al*., 1996). These include the gram-negative bacterium *S. typhimurium* AR3612, a *cysG metE* mutant, and the microalga *Euglena gracilis*. However, both *E. gracilis* (Helliwell *et al*., 2016) and *S. typhimurium* strain AR3612 (**Figure S4**) can grow on pseudocobalmin, indicating that they are unsuitable for distinguishing between benzimidazole and purinyl lower axial ligands. In contrast, both *C. reinhardtii* metE7 and *P. tricornutum* C9 are suitable candidates for developing a novel bioassay that would not only quantify cobalamin in a sample but could in principle also distinguish if pseudocobalamin, or other purine analogues, were present. To develop this, we chose to use the *C. reinhardtii* metE7 strain because, although *P. tricornutum* C9 has a lower vitamin requirement, it requires a much longer incubation time (7 days compared to 4 days) and the sensitivity meant that the transition between no growth and growth was more abrupt, so less easy to distinguish between different concentrations.

Since B_12_ is a cofactor for METH, as a first step it was essential to establish whether the presence of pseudocobalamin interfered with growth when both B_12_ vitamers were present. Accordingly, *C. reinhardtii* metE7 was grown on a range of cobalamin concentrations (5-325 pM) with or without the addition of 25 pM pseudocobalamin, and vice versa (**Figure 3**, **Figure S5**). When grown on cobalamin alone, the addition of DMB did not alter the growth of *C. reinhardtii* metE7 (**Figure 3a**). In contrast, when 25 pM pseudocobalamin was also present, DMB had a significant positive effect up to 81 pM cobalamin (red triangles), after which the growth potential was saturated (**Figure 3b**, Tukey post-hoc test all *p*<0.001 40 pM and below). In the reverse experiment, as expected, with pseudocobalamin alone growth was only observed when DMB was supplied (**Figure 3c**, blue triangles). When 25 pM cobalamin was also present (**Figure 3d**) then growth in the absence of DMB was supported, but only to the level of cobalamin provided (**Figure 3d**). From this experiment, it can be concluded that pseudocobalamin *per se* does not act as an antimetabolite, and that the growth of *C. reinhardtii* metE7 mutant is sensitive to very low levels of both this compound and cobalamin.

**Figure 3.**
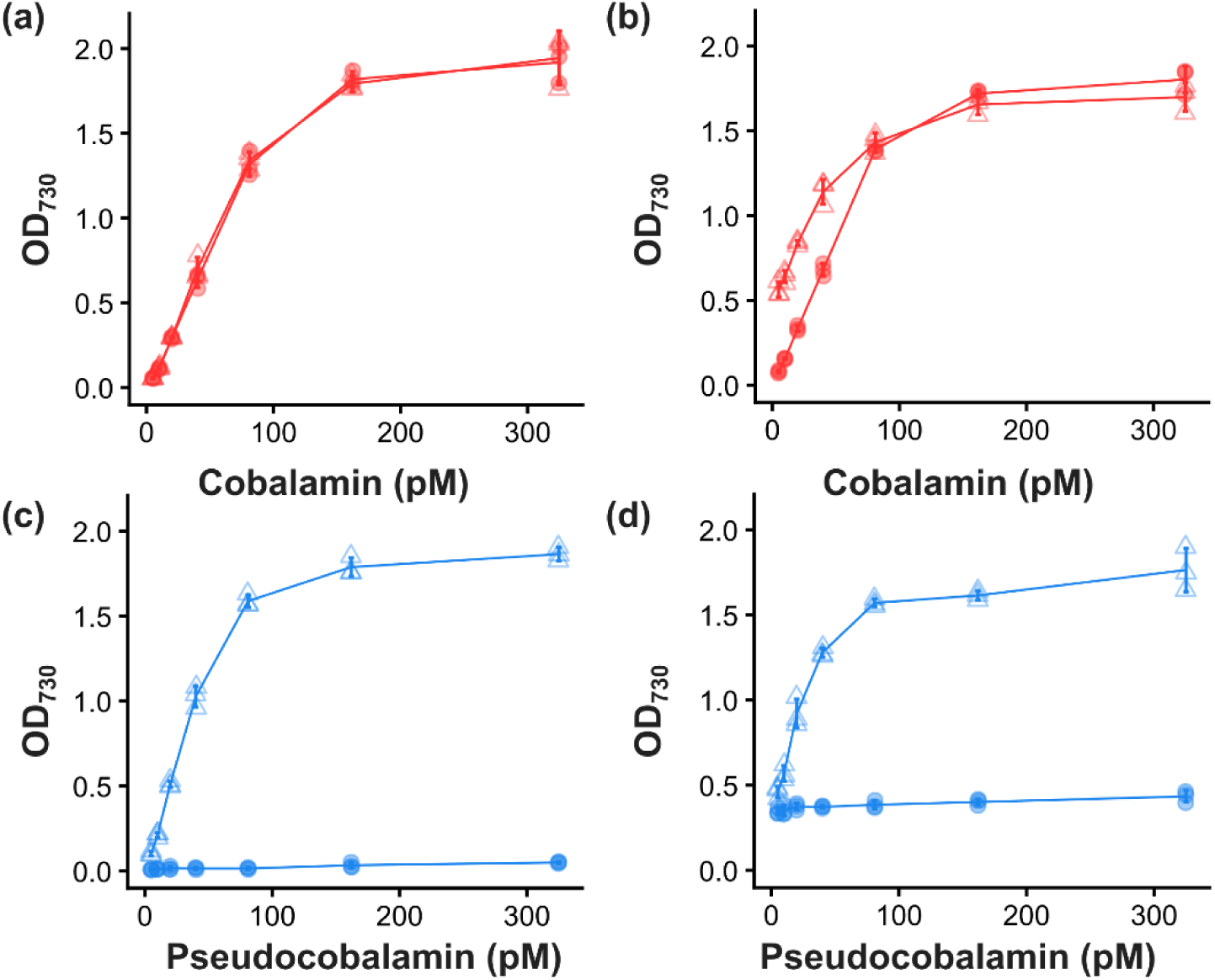
**Effect of B_12_ mixtures on the growth of *C. reinhardtii* metE7**. **a)** Growth on various concentrations (pM) of cobalamin (Cbl, shown in red) with (triangles) or without (circles) the addition of 1 µM DMB. **b**) the same experiment as in (a) but with the addition of 25 pM pseudocobalamin (PsCbl). **c)** growth at various concentrations of PsCbl (blue) with (triangles) or without (circles) addition of 1 µM DMB **d**) The same experiment as in (c) but with the addition of 25 pM Cbl. Optical density at 730 nm (OD_730_) as a proxy for growth, taken after 4 days of culturing. Line joins the average value with error bars, n=3 ±SD.

These growth data were used to generate a standard curve and fit to a 4-parameter logistic equation (Ritz *et al*., 2015) (**Figure S6, Methods S1, Table S4**), the parameters of which could then be used to compare the growth of *C. reinhardtii* metE7 on the various mixtures of B_12_ to the predicted values, which were in good agreement (**Figure S6**).

### CBA1 is necessary for the accumulation of both cobalamin and pseudocobalamin in two diverse microalgae

To gain insight into B_12_ uptake and potential benefits of the ability of microalgae to remodel, we investigated whether there was competition between the vitamers for uptake into the cell. The uptake of B_12_ by microalgae is less well characterised than other organisms, however recent progress has shown that CBA1 is crucial for the accumulation of cobalamin in both diatoms and *C. reinhardtii* (Bertrand *et al*., 2012; Sayer *et al*., 2024). We therefore investigated if it were also involved in pseudocobalamin uptake, taking advantage of previously generated *CBA1* knockout strains of both *C. reinhardtii* and *P. tricornutum* (Sayer *et al*., 2024). These mutants, together with their respective parental strains, were grown to the same growth stage, and the cell density normalised before the addition of a known amount of B_12_ vitamer for a set time. Samples were separated into cell and media fractions with the amount of B_12_ in each fraction quantified using the *C. reinhardtii* metE7 bioassay. **Figure 4a** shows the results for *C. reinhardtii CBA1* knockout strain, known as IM4, together with its parental strain UVM4 and a line of the mutant complemented with the wild-type CBA1 gene (IM4_comp). For UVM4 and IM4_comp, the majority of both forms of B_12_ was found in the cellular fraction after incubation, whereas when the mutant was incubated with either cobalamin or pseudocobalamin, all the B_12_ remained in the media fraction, indicating that a functional CBA1 was necessary for uptake of both vitamers. For the *P. tricornutum* lines, a similar uptake pattern was found (**Figure 4b**). In the bi-allelic knockout mutants, ΔCBA1-2 and ΔCBA1-3, there was no detection of either cobalamin or pseudocobalami in the cellular fraction, whereas the mono-allelic mutant (ΔCBA1-1) showed an intermediate phenotype, with both forms of B_12_ detected in the cell and media fractions. This suggests that the at least one functional *PtCBA1* copy is required for uptake of both cobalamin and pseudocobalamin.

**Figure 4.**
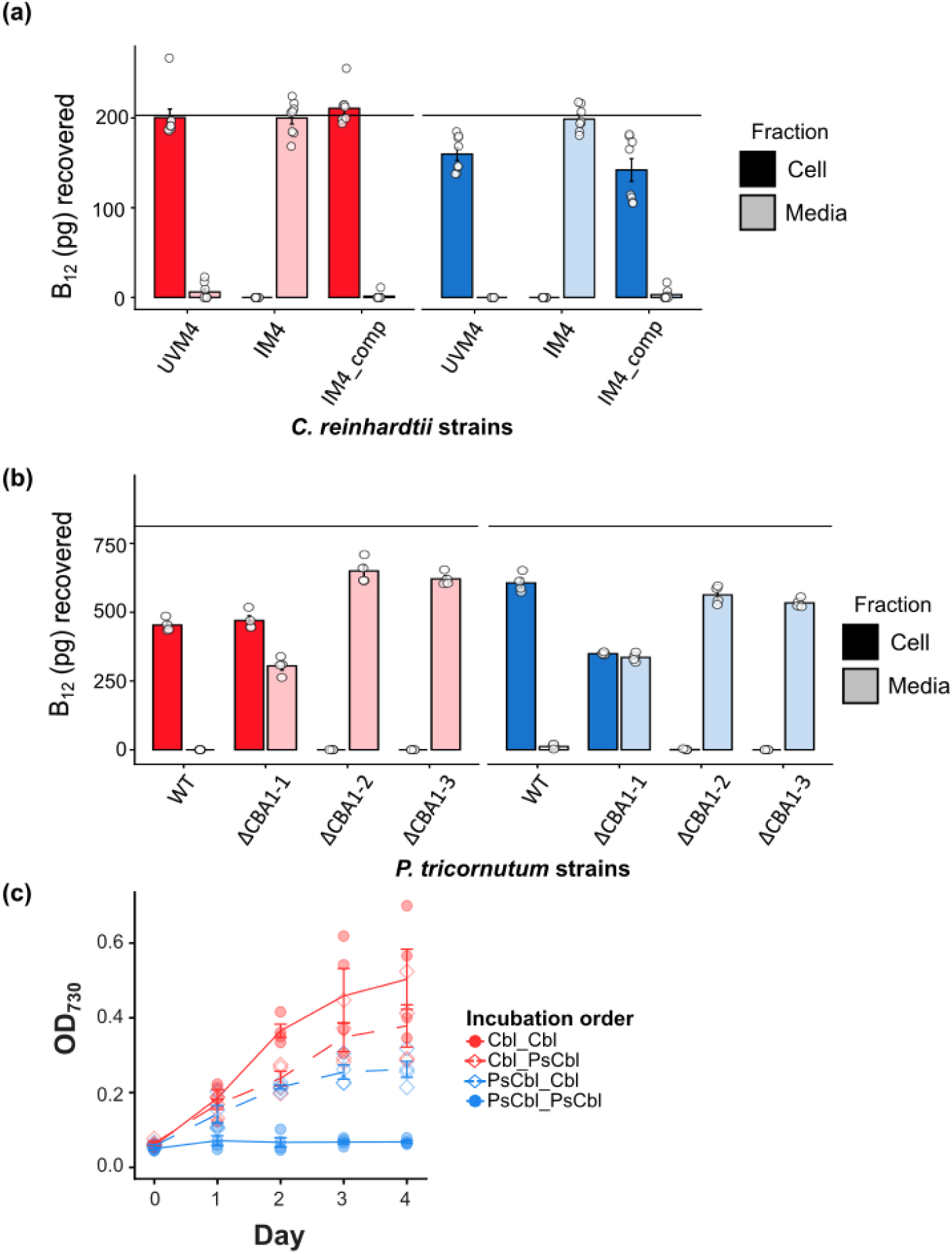
Uptake of cobalamin (Cbl) and pseudocobalamin (PsCbl) is via the same route in *C. reinhardtii* and *P. tricornutum*. **(a)** Cells of the *cba1-1* mutant (IM4) of *C. reinhardtii* (Sayer *et al*., 2024), together with its parental strain UVM4 and a strain complemented with the WT *CBA1* gene (IM4_comp) were incubated with TAP supplemented with 150 pM (∼200 g/ml) of either Cbl or PsCbl for 30 min. The amount of B_12_ recovered in the cell and media fractions was determined using the metE7 bioassay (material and methods). n=≥7 ±SE. **(b)** Cells of WT *P. tricornutum* and three independent CRISPR-Cas9 *PtCBA1* knockout lines (ΔCBA1) were incubated in modified F/2 supplemented with 600 pM (∼800 pg/ml) of either Cbl or PsCbl for 60 min, then the amount of B_12_ recovered in the cell and media fractions was determined, this was performed twice on the same samples with the average depicted. n=4 ±SE. The horizontal line indicates the approximate amount of B_12_ added to the samples, Cbl shown in red and PsCbl in blue. **(c)** Interference assay showing that Cbl and PsCbl are taken up by the same mechanism in *C. reinhardtii metE7*. Cells were incubated for 30 min with 1000 pM Cbl, followed by 30 min in 1000 pM PsCbl (red open diamond), cells were then washed, diluted and the OD_730_ (optical density at 730 nm) monitored for 4 days. Similar experiments were carried out for PsCbl then Cbl (blue open diamonds), Cbl then Cbl (red symbols) and PsCbl then PsCbl (blue symbols), n=4

We then carried out an uptake competition assay between the two forms of B_12_. *C. reinhardtii* metE7 cells were sequentially incubated with cobalamin for 30 min, washed and then incubated with pseudocobalamin for 30 min (**Figure 4c**, red open diamonds), or vice versa (blue open diamonds). Cells were washed again and then grown for 4 day in TAP media without any further B_12_ supplementation. As controls, cells were also incubated twice with cobalamin (red) or pseudocobalamin (blue). As expected, *C, reinhardtii* metE7 incubated twice with cobalamin reached the highest OD_730_, while those incubated twice with pseudocobalamin showed no growth. The cells incubated first with pseudocobalamin and then cobalamin reached an OD_730_ of 0.26 ± 0.04 (SD) compared to those incubated first with cobalamin which reached a higher OD _730_ of 0.38 ± 0.11 (SD), which while not statistically significant equates to a ∼1.5x higher density, and so is likely to be biologically relevant. This suggests that pseudocobalamin uptake interferes with that of cobalamin uptake, which in the absence of DMB is detrimental to the growth of the cells. The viability of the cells after the sequential incubations was tested by resupply of cobalamin to a dilution of the incubated cells, which all grew to the same extent (**Figure S7**). This indirect evidence suggests that cobalamin and pseudocobalamin are acquired by the same mechanism.

### *CrCOBT* is necessary for *C. reinhardtii* to remodel pseudocobalamin to enable growth

Several bacteria have been shown to remodel the lower axial ligand of different cobamides when DMB is present (Gray & Escalante-Semerena, 2007; Yi *et al*., 2012; Shelton *et al*., 2018). The enzymes responsible have been identified in the bacterium *Salmonella entrica* as CobT, CobS and CobC (Anderson *et al*., 2008). Investigation of the presence of *COBT* and *COBS* in a range of eukaryotic microalgae showed that both had a scattered distribution across the different phylogenetic groups, with *COBT* more common than *COBS* (Helliwell, 2017; Vancaester *et al*., 2020). *C. reinhardtii* encodes homologues of all three proteins (Helliwell *et al.,* 2016). To investigate this further, we used CRISPR/Cas9 gene-editing to disrupt the *COBT* gene in *C. reinhardtii.* We used the metE4 strain (Bunbury *et al*., 2020) since it was generated in the cell wall deficient UVM4 background, which makes transformation more efficient. The metE4 mutant behaves as the metE7 strain, in that it can grow in media supplemented with cobalamin or other benzimidazole analogues (**Figure S8**), but on pseudocobalamin or Factor A only when DMB is present (**Figure 5a)**. The gene editing design included a ssDNA repair template to insert a stop codon in *CrCOBT* and thus render it non-functional (see Materials and Methods & **Figure S2** for details). Transformants were picked into 96-well plates and several lines were found that were unable to grow with pseudocobalamin +DMB (**Figure S3a**). Four of the lines were taken, grown in TAP with cobalamin and genotyped by PCR of the target region of the COBT gene. In each case, the coding sequence had been edited to include a stop codon, so that it would no longer encode a functional COBT. We analysed one line, metE4_cobTD5, in more detail. **Figure 5d** shows that it was unable to grow on pseudocobalamin or Factor A in the presence of DMB, unlike the parental metE4 strain, confirming that the *COBT* gene is essential for remodelling activity.

**Figure 5.**
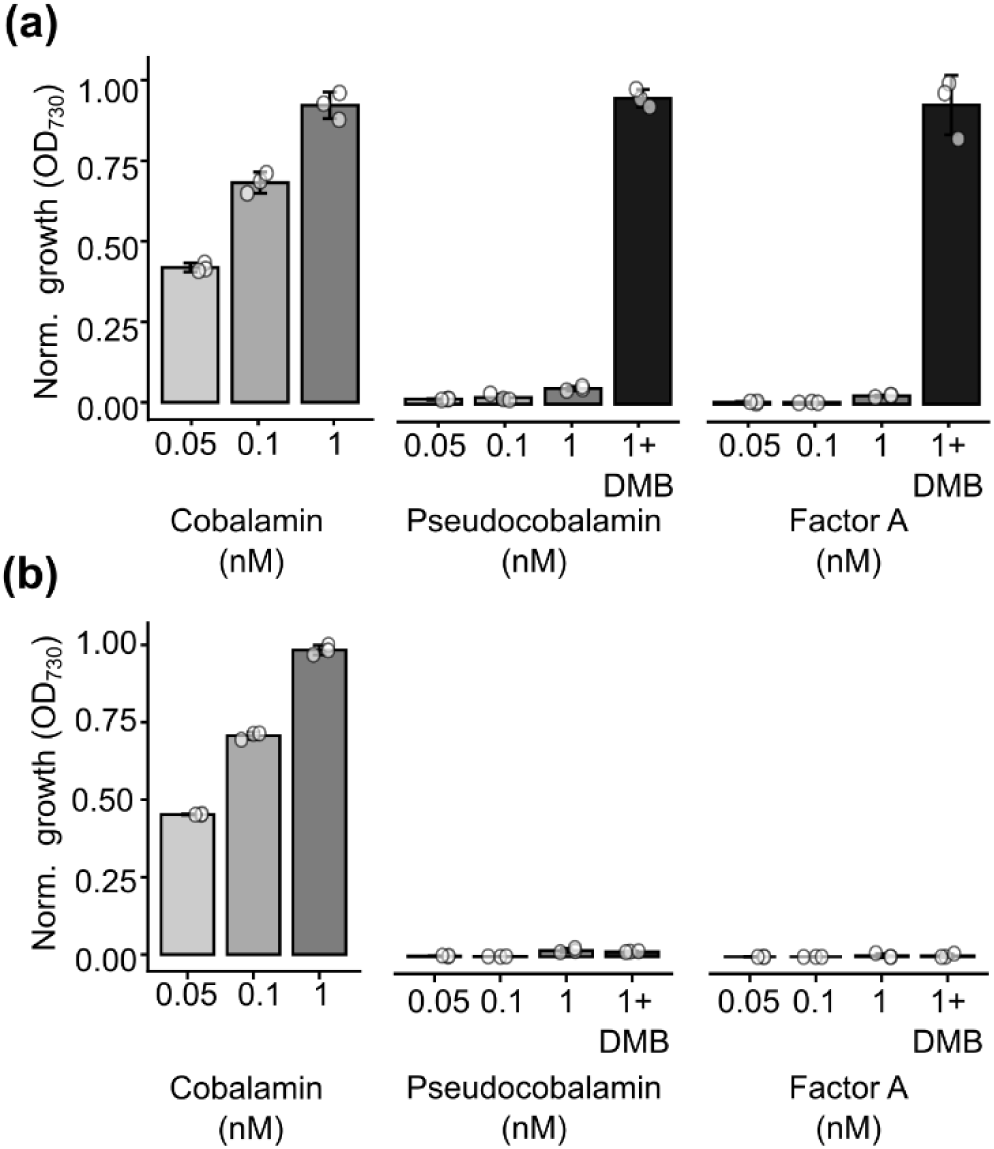
COBT is essential for remodelling of purinyl B_12_ analogues in *C. reinhardtii*. **a)** the growth of background strain *C. reinhardtii* metE4 on cobalamin, pseudocobalamin and Factor A, with and without the addition of 1 µM DMB. **b)** show the same experiments as **a**) but with the *C.reinhardtii* metE4 COBT knockout line, metE4_cobTD5. Both strains were grown in liquid media for 4-days. Growth measured using optical density at 730 nm and normalised by dividing all values by the highest optical density achieved by each strain. (n=3, mean ±SD).

### *Chlorella vulgaris* can accumulate pseudocobalamin but cannot remodel it

*C. vulgaris*, and *Chlorella* more generally, is often cited as a future food and good source of vitamin B_12_ for human consumption (Kittaka-Katsura *et al*., 2002; Bito *et al*., 2016; Wells *et al*., 2017; Parodi *et al*., 2018; Durdakova *et al*., 2024). A wide range of B_12_ concentrations have been detected in *Chlorella* biomass marketed as food supplements (Kittaka-Katsura *et al*., 2002; Bito *et al*., 2016), however details of the algal cultivation are often lacking, including the species and strain and whether the system is open or closed (Bito *et al*., 2016). Since B_12_ is not biosynthesised by the alga, the vitamin must have been acquired from the cultivation environment. Moreover, as pseudocobalamin and other purinyl cobamides are less bioavailable to humans (Stupperich & Nexø, 1991; Fedosov *et al*., 2007; Sokolovskaya *et al*., 2021), it is important to characterise the form(s) of B_12_ found in *Chlorella*, including its ability to take up different vitamers. There are no reports of B_12_-dependence in the *Chlorella* genus (Provasoli, 1958; Provasoli & Carlucci, 1974), and inspection of the published genome for *Chlorella variabilis* confirm the presence of a complete *METE* gene (Helliwell et al., 2011; 2017). One study assessed the ability of 9 strains, including 3 strains of *Chlorella pyrenoidosa*, 3 of *C. vulgaris*, *Chorella ellipsoidea* C-27, *Chlorella miniata* C-143 and *Chlorella zopfingiensis* C-111, to accumulate vitamin B_12_ over several days, and found considerable variability (Maruyama *et al*.,1989). We therefore decided to carry out a more systematic investigation, using the widely-studied *C. vulgaris* strain 211/11B, to determine its ability to accumulate cobalamin and pseudocobalamin. Cells were incubated with either 150 pM cobalamin or pseudocobalamin, followed by separation of the cell and media fractions by centrifugation. B_12_ uptake was then assayed by measuring B_12_ levels in the two fractions, using the *C. reinhardtii* metE7 bioassay.

As with the other algae tested, *C. vulgaris* 211/11B could accumulate both cobalamin and pseudocobalamin, with the majority of the added B_12_ being recovered from the cellular fraction after 30 minutes (**Figure 6a**). However, there was a difference in the propensity of the cells to accumulate cobalamin (180.5 pg ±7.0 SE) compared to pseudocobalamin (117.4 pg ±3.0 SE) (t-test, *p*<0.01). When different timepoints were taken across the 30 minute incubation to assess the uptake dynamics, within 2 minutes (120 secs) it was possible to detect B_12_ in the cellular fraction for both vitamers (**Figure 6b**), with ∼55% of the added cobalamin and ∼43% of pseudocobalamin after 5 minutes. However, uptake of pseudocobalamin then plateaued, whereas for cobalamin the amount continues to increase until ∼73% of the total B_12_ was detected in the cell fraction (**Figure 6b**).

**Figure 6.**
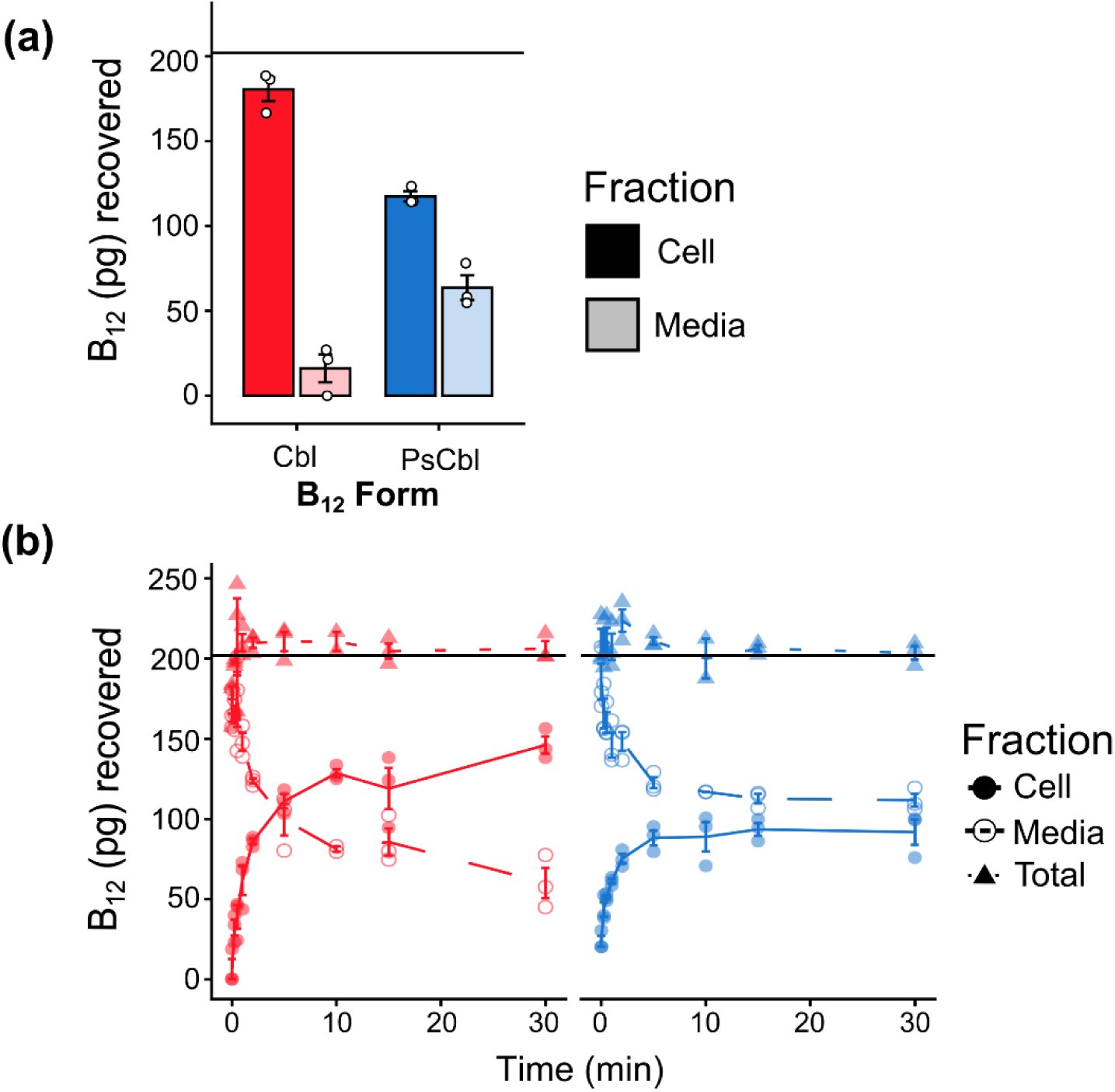
***Chlorella vulgaris* takes up both cobalamin (Cbl) and pseudocobalamin (PsCbl)**. **a)** *C. vulgaris* 211/11B cells were incubated for 30 minutes with 150 pM B_12_ either in the form of Cbl (203 pg/ml) or PsCbl (201 pg/ml) and then processed as detailed in Methods. n=3 ±SE. **b)** Time course of B_12_ uptake by *C. vulgaris* 211/11B over 30 minutes. The horizontal line indicates the approximate total amount added to the samples, red indicates Cbl and blue indicates PsCbl. n=3* ±SE. B_12_ (pg) recovered by the B_12_-dependent *C. reinhardtii* metE7 bioassay as detailed in the methods. *See methods

Despite the higher uptake of cobalamin over pseudocobalamin, *C. vulgaris* still accumulated approximately half of the pseudocobalamin provided. Therefore, we wanted to investigate whether *C. vulgaris* was capable of remodelling pseudocobalamin to cobalamin. A previous BLASTP search of the *C. variabilis* genome had not detected genes for COBS, T or C (Helliwell, 2017), whereas *C. sorokiniana* possesses homologues for the three putative remodelling genes (Arriola *et al*., 2018; Durdakova *et al*., 2022). Phylogenetic analysis inferred from rRNA sequence data suggests that *C. variabilis* is more closely related to *C. vulgaris* than *C. sorokiniana* (Bock *et al*., 2011). A BLASTP search of *Chlorella vulgaris* (taxid:3077) using protein sequences of *COBT* from various taxa (*S. typhimurium*-NP_460961.1, *C. reinhardtii*-XP_042918919.1, *P. tricornutum*-XP_002184872.1 and *C. sorokiniana*-PRW60684.1) returned no significant hits.

The bioinformatics analysis suggests that *C. vulgaris* is unable to remodel, but to be certain it was necessary to assess this trait experimentally. To do this, we took advantage of a naturally-occurring B_12_-dependent alga, *Lobomonas rostrata*, which is incapable of remodelling pseudocobalamin (Helliwell *et al*., 2016) and so can grow only on supplemented cobalamin. By using this as an indicator strain, we could determine if extracts from other algal strains supplied with pseudocobalamin+DMB contained cobalamin. Cultures of *C. vulgaris* 211/11B and *C. reinhardtii* WT-12 were grown in TAP medium for 2 days before addition of 250 pM pseudocobalamin + 5 nM DMB. Growth was continued for a further 6 days, with 1 ml samples taken daily. The cells were recovered by centrifugation, boiled and the soluble extracts used to supplement the growth of the indicator strain *L. rostrata*. As a control for any negative effects of cell contents on the indicator strain, *C. vulgaris* 211/11B and *C. reinhardtii* WT-12 were grown in the same way as above but with the addition of 250 pM cobalamin. **Figure 7** shows the growth of *L. rostrata* supplemented with cell extracts from the different treatments at day 4, but similar trends were seen with all timepoints (**Figure S9**). Extracts of *C. reinhardtii* WT-12 grown either with cobalamin (red symbols) or pseudocobalamin+DMB (blue symbols) were able to support *L. rostrata* growth (**Figure 7a**). This indicates that pseudocobalamin remodelled by *C. reinhardtii* WT-12 was bioavailable to the indicator strain, although the growth achieved was less than when using the cobalamin extracts. In contrast, only those cellular extracts of *C. vulgaris* supplemented with cobalamin supported *L. rostrata* growth (**Figure 7b**). This experimental evidence supports the conclusion that *C. vulgaris* cannot remodel pseudocobalamin.

**Figure 7.**
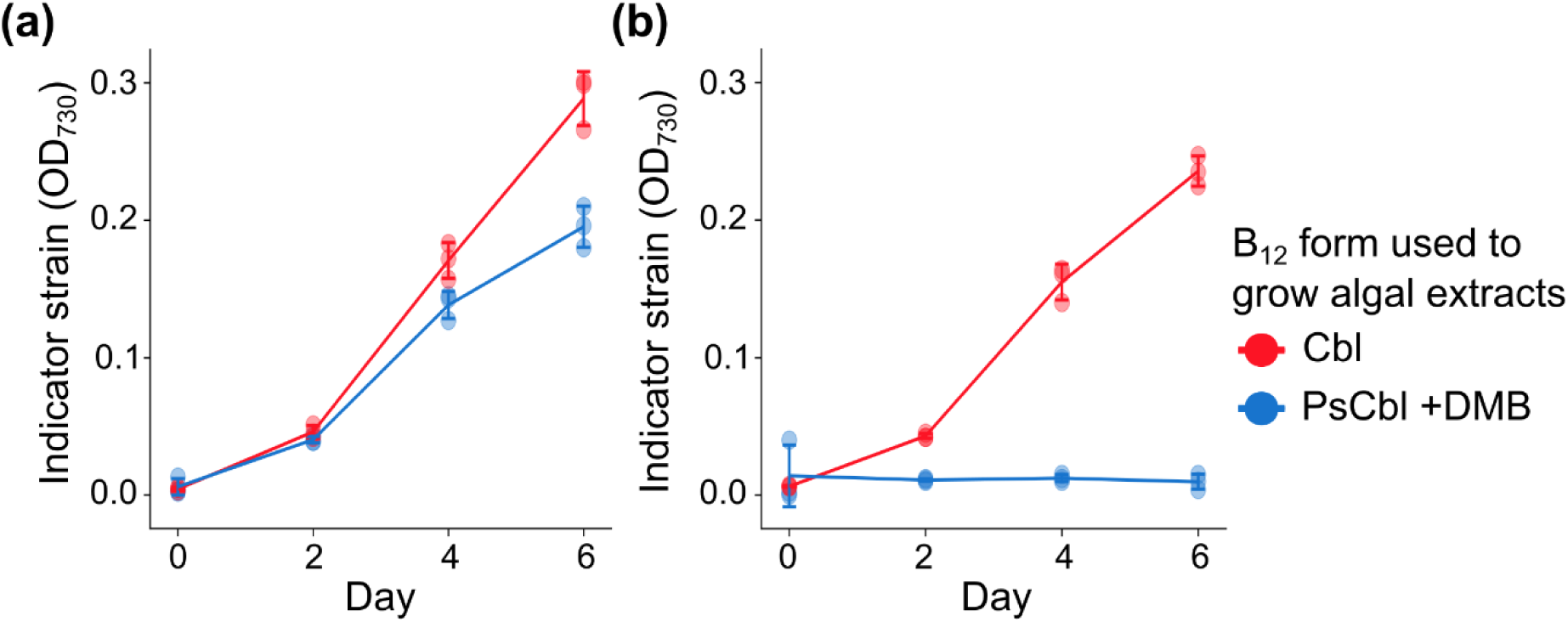
Extracts of *C. vulgaris* grown with pseudocobalmin and DMB cannot support the growth of a non-remodelling B_12_-dependent alga. Cellular extracts of *C. reinhardtii* (**a**) and *C. vulgaris* (**b**) were prepared by growing each alga either on cobalamin (red) or pseudocobalamin supplemented with DMB (blue), see methods for details. These extracts were then used to supplement the medium of the indicator strain *L. rostrata,* with the growth monitored using optical density at 730 nm (OD_730_) over 6 days. The *C. vulgaris* extract was unable to support *L. rostrata* growth, showing that the former was not able to remodel, unlike *C. reinhardtii*. n=3 ±SD

## Discussion

This work builds on earlier studies of algal B_12_ metabolism, uptake and remodelling to provide a more detailed picture of these crucial processes. Firstly, we have expanded the known bioavailability of different benzimidazole vitamers of B_12_ for the B_12_-dependent *C. reinhardtii* metE7 mutant and shown it is able to remodel Factor A as well as pseudocobalamin (**Figure 2a & b)**. Identical results were obtained with B_12_-dependent *P. tricornutum* C9 strain that we generated by gene-editing (**Figure 2c & d**), although lower levels of B_12_ are required to support growth of this diatom than for the green alga *C. reinhardtii*. Second, we exploited this remodelling phenotype to develop a novel microbiological assay using *C. reinhardtii* metE7, which allows the quantification of B_12_ at pM concentrations and which is able to distinguish between B_12_ vitamers with benzimidazole and adenine lower axial ligands in samples (**Figure 3, Figure S6**). We also established that CBA1, so far the only known protein involved in algal B_12_ uptake (Bertrand *et al*., 2012; Sayer *et al*., 2024) is also necessary for the accumulation of pseudocobalamin in these two highly distinct algal taxa (**Figure 4**). Recent findings also suggest the presence of CBA1 homologues in some vascular plants and hornworts; many hornworts contain cyanobacterial symbionts that may produce pseudocobalamin (Dorrell *et al*., 2024), a potential interaction that could be further investigated with this novel microbiological assay.

Given the taxonomic distance between *C. reinhardtii* and *P. tricornutum* the similarity in the metabolism and uptake routes of B_12_ in the two species is remarkable, especially since other more closely-related species do not share these traits. Vitamin B_12_ is known to limit the growth of phytoplankton more generally, particularly in polar regions (Bertrand *et al*., 2007; Koch *et al*., 2011; Browning *et al*., 2017), therefore the ability to utilise other corrinoid compounds through scavenging and remodelling would provide a competitive advantage to those species that possess the capacity. The presence of DMB has been quantified in soil and freshwater environments (Crofts *et al*., 2014), and more recently in marine waters (Bruns *et al*., 2022; Bannon *et al*., 2025). One study of the Northwest Atlantic found 6.9 ± 5.6 pM DMB in the dissolved fraction across most of the sampled transect, with seasonal dynamics detected in the particulate fraction, which were suggested to be associated with differences in the importance of cobamide remodelling during the autumn bloom compared to spring (Bannon *et al*., 2025). There is also evidence for the presence of α-ribazole, a DMB intermediate in the exometabolomes of marine bacteria (Johnson *et al*., 2016; Wienhausen *et al*., 2017) and as a photodegradation product of hydroxy-cobalamin (Bannon *et al*., 2024). One study showed that supplementation with cobalamin or α-ribazole altered the community dynamics of prokaryotes and heterotrophic protists present (Wienhausen *et al*., 2022). The presence of the other analogues used in this study have not been quantified in the marine environment, however they have been found in samples from several microbial communities, such as those isolated from soil (Men *et al*., 2015), human faeces (Allen & Stabler, 2008), and the rumen of cows (Girard *et al*., 2009), with Factor A being particularly predominant in human-associated samples (Allen & Stabler, 2008). It is therefore likely that in areas which gain nutrient inputs from wastewater and agricultural runoff, the remodelling of different forms of B_12_ provides a valuable resource to phytoplankton, shaping community dynamics as has been suggested for prokaryotes (Shelton *et al*., 2018).

With regards to how microalgae remodel unsuitable corrinoids to more bioavailable forms we have confirmed that *CrCOBT* is essential for remodelling in *C. reinhardtii* (**Figure 5**). By investigating the function of CobT in several diverse bacteria, it has been shown that CobT specificity determines the range of cobamides that are synthesised, functioning as a protective measure to ensure that the corrinoid produced supports the metabolism of the specific bacterium (Crofts *et al*., 2013). While the biochemical mechanism of how COBT functions in microalgae is still not known, several microalgae contain homologues of *COBT*, *COBS* and *COBC* (Helliwell, 2017; Vancaester *et al*., 2020). In the bacterium *Vibrio cholerae*, CobS was shown to be necessary for remodelling of pseudocobalamin to cobalamin (Ma *et al*., 2020). We also attempted to generate a *COBS* knockout mutant in *C. reinhardtii* using the same phenotypic screen as for *COBT*, namely the inability to grow on pseudocobalamin supplemented with DMB; however no lines were obtained with this phenotype. While this could be due to low transformation or editing efficiency, it might also suggest some functional redundancy in the biochemical mechanism of remodelling. Further, given the slight prevalence of *COBT* over *COBS* in the surveyed microalgal genomes (Helliwell *et al*., 2016; Helliwell, 2017; Vancaester *et al*., 2020) it may not be essential to the process. This warrants greater investigation as more genomes become available. In this context, the use of the non-remodelling *L. rostrata* as an indicator strain (**Figure 7**) was crucial in providing experimental support for the genetic approach. Another route to remodel pseudocobalamin to cobalamin, found in certain archaea and bacteria and characterised in the bacterium *Rhodobacter sphaeroides*, involves the aminohydolase CbiZ (Escalante-Semerena, 2007; Gray & Escalante-Semerena, 2009). No potential CbiZ homologues were found in *C. reinhardtii* (Helliwell *et al*., 2016), but it has been found in several diatom genomes, suggested to be the result of horizontal gene transfer from bacteria (Vancaester *et al*., 2020). That analysis did not identify *CbiZ*, or indeed any of the other final synthesis genes, in *P. tricornutum* (Vancaester *et al*., 2020), suggesting that CobT could be essential for remodelling in the diatom as well. Not only does this work provide greater insight into B_12_ metabolism in microalgae, the *CrCOBT* knockout line could be used as an additional strain in the bioassay detailed here, giving some indication of the presence or absence of DMB in unknown samples (**Figure 5**).

Finally, we have applied the bioassay approach and reporter strains we generated to investigate the capability of *C. vulgaris* to take up B_12_, adding to evidence from algal food supplements that *C. vulgaris* is capable of accumulating the compound (Kittaka-Katsura *et al*., 2002; Bito *et al*., 2016). There was a slight preference for cobalamin over pseudocobalamin (**Figure 6b**), but the latter was still efficiently taken into the cell. As this is of relevance to human health with *Chlorella* proposed as a dietary supplement (Tokuşoglu & Ünal, 2003; Merchant *et al*., 2015; Parodi *et al*., 2018), we investigated if *C. vulgaris* 211-11B was capable of remodelling pseudocobalamin to the more bioavailable form for humans (Fedosov *et al*., 2007; Sokolovskaya *et al*., 2021) and found that it did not have this capability (**Figure 7**), highlighting the need to consider the species of *Chlorella* selected when producing human diet supplements. One study of growing *C. vulgaris* in open-ponds detected methyl-cobalamin with tandem mass spectrometry (MS/MS), at approximately 28 µg/100g dry weight (Kumudha *et al*., 2015) suggesting that suitable microbial communities can colonise *C. vulgaris* ponds for human diet supplements. However, given the unregulated nature of health supplement production, the fact that at least one species of *Chlorella* is not capable of remodelling pseudocobalamin warrants more consideration. Along with the more applied investigations, the novel microbiological assay will enable us to address fundamental questions of metabolism and community interactions in both marine and freshwater microalgae that centre on the enigmatic vitamin, B_12_.

## Acknowledgements

We are grateful to Prof Martin Warren (University of Kent, UK) and Prof Dr Bernhard Kräutler (University of Innsbruck, Austria) for the B_12_ vitamers. The plasmids used for CRISPR/Cas9 editing of *P. tricornutum* were gifts from Dr Amanda Hopes and Professor Thomas Mock (UEA, Norwich, UK) and are also available from Addgene. This work was supported by MELiSSA Foundation PS POMP programme Doctoral Award to MPD & AGS, which provided a PhD studentship to ELH; the UK’s Biotechnology and Biological Sciences Research Council (BBSRC) for Doctoral Training grant (no. BB/M011194/1) to FB, APS, ML-P and AGS; grant no. BB/M018180/1 to P.M. and A.G.S; grant nos. BB/L002957/1 and BB/R021694/1 to KG and AGS.

## Author contributions

AGS, MPD and PM conceived and designed the research and obtained funding; ELH, FB, TS, APS, MLP, KP, KG, PM, MPD and AGS planned the experimental work; ELH, FB, TS, APS, MLP, KP and KG performed the experiments and data analysis. ELH and AGS wrote the manuscript with contributions from all authors. All authors reviewed and accepted the submitted manuscript.

## Data availability

All raw data, query sequences and scripts to generate the figures in this paper can be found at https://github.com/ellenharrison/Cellular_basis_Harrison-et-al_2026.git

## Supporting Information

**Table S1** Strains used in this work, and maintenance and experimental conditions.

**Table S2** Details of the cell density and concentration of B_12_ used for the uptake experiments.

**Table S3** Details of oligonucleotides used in this study.

**Table S4** The parameters used to relate the growth of bioassay strain to the concentration of B_12_.

**Fig. S1** Genotyping and characterisation of CRISPR/Cas9 PtMETE mutants

**Fig. S2** Strategy for the generation of CrCOBT mutant

**Fig. S3** Identifying CrCOBT mutant

**Fig. S4** The growth of *Salmonella typhimurium* on mixtures of B_12_

**Fig. S5** Effect of the presence of cobalamin and pseudocobalamin on the growth of *C. reinhardtii* metE7

**Fig. S6** Example of standard curve from *C. reinhardtii* metE7 bioassay

**Fig. S7** Confirmation of *C. reinhardtii* metE7 cell viability after sequential incubation with B_12_ analogues

**Fig. S8** *C. reinhardtii* metE4 and CrCOBT knockout line growth on 3 other benzimidazole B_12_ analogues

**Fig. S9** The growth of indicator strain *L. rostrata* on extracts of *C. vulgaris* or *C. reinhardtii* which in turn were grown with cobalamin or pseudocobalamin+DMB

**Methods S1** Equations necessary for the bioassay

## References

Allen RH, Stabler SP. 2008. Identification and quantitation of cobalamin and cobalamin analogues in human feces. American Journal of Clinical Nutrition 87: 1324–1335.

Anderson BB. 1964. Investigations Into the Euglena method for the assay of the vitamin B12. Journal of Clinical Pathology 17: 14–26.

Anderson PJ, Lango J, Carkeet C, Britten A, Kräutler B, Hammock BD, Roth JR. 2008. One pathway can incorporate either adenine or dimethylbenzimidazole as an α-axial ligand of B12 cofactors in *Salmonella enterica*. Journal of Bacteriology 190: 1160–1171.

Arriola MB, Velmurugan N, Zhang Y, Plunkett MH, Hondzo H, Barney BM. 2018. Genome sequences of *Chlorella sorokiniana* UTEX 1602 and *Micractinium conductrix* SAG 241.80: implications to maltose excretion by a green alga. Plant Journal 93: 566–586.

Bannon CC, Mudge EM, Bertrand EM. 2024. Shedding light on cobalamin photodegradation in the ocean. Limnology And Oceanography Letters 9: 135–144.

Bannon C, White PL, Rowland E, More KJ, Gleason A, Roberts M, Devred E, Beazley L, LaRoche J, Bertrand EM. 2025. Seasonal patterns in B-vitamins and cobalamin co-limitation in the Northwest Atlantic. Limnology and Oceanography 70: 3370–3385.

Becker EW. 2007. Micro-algae as a source of protein. Biotechnology Advances 25: 207–210.

Bertrand EM, Allen AE, Dupont CL, Norden-Krichmar TM, Bai J, Valas RE, Saito MA. 2012. Influence of cobalamin scarcity on diatom molecular physiology and identification of a cobalamin acquisition protein. Proceedings of the National Academy of Sciences USA 109: E1762–E1771.

Bertrand EM, Saito MA, Rose JM, Riesselman CR, Lohan MC, Noble AE, Lee PA, DiTullio GR. 2007. Vitamin B_12_ and iron colimitation of phytoplankton growth in the Ross Sea. Limnology and Oceanography 52: 1079–1093.

Bito T, Bito M, Asai Y, Takenaka S, Yabuta Y, Tago K, Ohnishi M, Mizoguchi T, Watanabe F. 2016. Characterization and Quantitation of Vitamin B _12_ Compounds in Various *Chlorella* Supplements. Journal of Agricultural and Food Chemistry 64: 8516–8524.

Bock C, Krienitz L, Pröschold T. 2011. Taxonomic reassessment of the genus *Chlorella* (Trebouxiophyceae) using molecular signatures (barcodes), including description of seven new species. Fottea 11: 293–312.

Bonnet S, Tovar-Sánchez A, Panzeca C, Duarte CM, Ortega-Retuerta E, Sañudo-Wilhelmy SA. 2013. Geographical gradients of dissolved vitamin B12 in the Mediterranean Sea. Frontiers in Microbiology 4: 1–10.

Browning TJ, Achterberg EP, Rapp I, Engel A, Bertrand EM, Tagliabue A, Moore CM. 2017. Nutrient co-limitation at the boundary of an oceanic gyre. Nature 551: 242–246.

Browning TJ, Rapp I, Schlosser C, Gledhill M, Achterberg EP, Bracher A, Le Moigne FAC. 2018. Influence of iron, cobalt, and vitamin B12 supply on phytoplankton growth in the Tropical East Pacific during the 2015 El Niño. Geophysical Research Letters 45: 6150–6159.

Bruns S, Wienhausen G, Scholz-Böttcher B, Wilkes H. 2022. Simultaneous quantification of all B vitamins and selected biosynthetic precursors in seawater and bacteria by means of different mass spectrometric approaches. Analytical and Bioanalytical Chemistry 414: 7839– 7854.

Bunbury F, Helliwell KE, Mehrshahi P, Davey MP, Salmon DL, Holzer A, Smirnoff N, Smith AG. 2020. Responses of a newly evolved auxotroph of *Chlamydomonas* to B12 deprivation. Plant Physiology 183: 167–178.

Cavari B, Grossowicz N. 1977. Seasonal Distribution of Vitamin B12 in Lake Kinneret. Applied and Environmental Microbiology 34:120-124.

Chimento DP, Mohanty AK, Kadner RJ, Wiener MC. 2003. Substrate-induced transmembrane signaling in the cobalamin transporter BtuB. Nature Structural Biology 10: 394–401.

Cole MB, Augustin MA, Robertson MJ, Manners JM. 2018. The science of food security. npj Science of Food 2: 1–8.

Croft MT, Lawrence AD, Raux-Deery E, Warren MJ, Smith AG. 2005. Algae acquire vitamin B12 through a symbiotic relationship with bacteria. Nature 438: 90–93.

Crofts TS, Men Y, Alvarez-Cohen L, Taga ME. 2014. A bioassay for the detection of benzimidazoles reveals their presence in a range of environmental samples. Frontiers in Microbiology 5: 1–12.

Crofts TS, Seth EC, Hazra AB, Taga ME. 2013. Cobamide structure depends on both lower ligand availability and CobT substrate specificity. Chemistry and Biology 20: 1265–1274.

Daisley KW. 1969. Monthly survey of vitamin B12 concentrations in some waters of the English Lake District. Limnology and Oceanography 14: 224–228.

Degnan PH, Barry NA, Mok KC, Taga ME, Goodman AL. 2014. Human gut microbes use multiple transporters to distinguish vitamin B12 analogs and compete in the gut. Cell Host and Microbe 15: 47–57.

Dorrell RG, Nef C, Altan-Ochir S, Bowler C, Smith AG. 2024. Presence of vitamin B_12_ metabolism in the last common ancestor of land plants. Philosophical Transactions of the Royal Society B: Biological Sciences 379: 20230354.

Durdakova M, Kolackova M, Janova A, Krystofova O, Adam V, Huska D, Durdakova M, Kolackova M, Janova A, Krystofova O, et al. 2022. Microalgae / cyanobacteria: the potential green future of vitamin B12 production. Critical Reviews in Food Science and Nutrition 64: 3091–3102.

Durdakova M, Kolackova M, Ridoskova A, Cernei N, Pavelicova K, Urbis P, Richtera L, Pelcova P, Adam V, Huska D. 2024. Exploring the potential nutritional benefits of *Arthrospira maxim*a and *Chlorella vulgaris*: A focus on vitamin B12, amino acids, and micronutrients. Food Chemistry doi: 10.1016/j.foodchem.2024.139434.

Escalante-Semerena JC. 2007. Conversion of cobinamide into adenosylcobamide in bacteria and archaea. Journal of Bacteriology 189: 4555–4560.

FAO. Fisheries and Aquaculture. 2008. A review on culture, production and use of Spirulina as food for humans and feeds for domestic animals and fish. Circular No. 1034.

Fedosov SN, Fedosova NU, Kräutler B, Nexø E, Petersen TE. 2007. Mechanisms of discrimination between cobalamins and their natural analogues during their binding to the specific B12-transporting proteins. Biochemistry 46: 6446–6458.

Ferenczi A, Chew YP, Kroll E, von Koppenfels C, Hudson A, Molnar A. 2021. Mechanistic and genetic basis of single-strand templated repair at Cas12a-induced DNA breaks in *Chlamydomonas reinhardtii*. Nature Communications 12: 6751.

Ford JE, Hutner SH. 1955. Role of vitamin B12 in the metabolism of microorganisms. Vitamins and Hormones 13: 101–136.

Girard CL, Santschi DE, Stabler SP, Allen RH. 2009. Apparent ruminal synthesis and intestinal disappearance of vitamin B12 and its analogs in dairy cows. Journal of Dairy Science 92: 4524–4529.

Gray MJ, Escalante-Semerena JC. 2007. Single-enzyme conversion of FMNH2 to 5,6-dimethylbenzimidazole, the lower ligand of B12. Proceedings of the National Academy of Sciences USA 104: 2921–2926.

Gray MJ, Escalante-Semerena JC. 2009. The cobinamide amidohydrolase (cobyric acid-forming) CbiZ enzyme: A critical activity of the cobamide remodelling system of *Rhodobacter sphaeroides*. Molecular Microbiology 74: 1198–1210.

Heal KR, Qin W, Ribalet F, Bertagnolli AD, Coyote-Maestas W, Hmelo LR, Moffett JW, Devol AH, Armbrust EV, Stahl DA, et al. 2017. Two distinct pools of B12 analogs reveal community interdependencies in the ocean. Proceedings of the National Academy of Sciences USA 114: 364–369.

Helliwell KE. 2017. The roles of B vitamins in phytoplankton nutrition: new perspectives and prospects. New Phytologist 216: 62–68.

Helliwell KE, Collins S, Kazamia E, Purton S, Wheeler GL, Smith AG. 2015. Fundamental shift in vitamin B12 eco-physiology of a model alga demonstrated by experimental evolution. The ISME Journal 9: 1446–1455.

Helliwell KE, Lawrence AD, Holzer A, Kudahl UJ, Sasso S, Kräutler B, Scanlan DJ, Warren MJ, Smith AG. 2016. Cyanobacteria and eukaryotic algae use different chemical variants of vitamin B12. Current Biology 26: 999–1008.

Helliwell KE, Wheeler GL, Leptos KC, Goldstein RE, Smith AG. 2011. Insights into the evolution of vitamin B 12 auxotrophy from sequenced algal genomes. Molecular Biology and Evolution 28: 2921–2933.

Hutner SH, Bach MK, Ross GIM. 1956. A sugar-containing basal medium for vitamin B12 assay with Euglena: application to body fluids. The Journal of Protozoology 3: 101–112.

Johnson WM, Kido Soule MC, Kujawinski EB. 2016. Evidence for quorum sensing and differential metabolite production by a marine bacterium in response to DMSP. ISME Journal 10: 2304–2316.

Kassambara A. 2025. ggpubr: ‘ggplot2’ based publication-ready plots. R package version 0.6.3.999. https://CRAN.R-project.org/package=ggpubr.

Kelleher BP, Broin SDO. 1991. Microbiological assay for vitamin B12 performed in 96-well microtitre plates. Journal of Clinical Pathology 44: 592–595.

Keller S, Kunze C, Bommer M, Paetz C, Menezes RC, Svatoš A, Dobbek H, Schubert T. 2018. Selective utilization of benzimidazolylnorcobamides as cofactors by the tetrachloroethene reductive dehalogenase of *Sulfurospirillum multivorans*. Journal of Bacteriology 200: e00584-17.

Kennedy KJ, Taga ME. 2020. Cobamides. Current Biology 30: R55–R56.

Kittaka-Katsura H, Fujita T, Watanabe F, Nakano Y. 2002. Purification and Characterization of a Corrinoid Compound from *Chlorella* Tablets as an Algal Health Food. Journal of Agricultural and Food Chemistry 50: 4994–4997.

Koch F, Marcoval MA, Panzeca C, Bruland KW, Sañudo-Wilhelmy SA, Gobler CJ. 2011. The effect of vitamin B12 on phytoplankton growth and community structure in the Gulf of Alaska. Limnology and Oceanography 56: 1023–1034.

Kumudha A, Selvakumar S, Dilshad P, Vaidyanathan G, Thakur MS, Sarada R. 2015. Methylcobalamin – A form of vitamin B12 identified and characterised in *Chlorella vulgaris*. Food Chemistry 170: 316–320.

Lin S, Hu Z, Song X, Gobler CJ, Tang YZ. 2022. Vitamin B12-auxotrophy in dinoflagellates caused by incomplete or absent cobalamin-independent methionine synthase genes (metE). Fundamental Research 2: 727-737.

Llavero-Pasquina M, Geisler K, Holzer A, Mehrshahi P, Mendoza-Ochoa GI, Newsad SA, Davey MP, Smith AG. 2022. Thiamine metabolism genes in diatoms are not regulated by thiamine despite the presence of predicted riboswitches. New Phytologist 235: 1853–1867.

Ma AT, Tyrell B, Beld J. 2020. Specificity of cobamide remodeling, uptake and utilization in Vibrio cholerae. Molecular Microbiology 113: 89–102.

Martens JH, Barg H, Warren M, Jahn D. 2002. Microbial production of vitamin B12. Applied Microbiology and Biotechnology 58: 275–285.

Maruyama I, Ando YH, Maeda T, Hirayama K. 1989. Uptake of vitamin B12 by various strains of unicellular algae *Chlorella*. Nippon Suisan Gakkaishi 55: 1785–1790.

Men Y, Seth EC, Yi S, Crofts TS, Allen RH, Taga ME, Alvarez-Cohen L. 2015. Identification of specific corrinoids reveals corrinoid modification in dechlorinating microbial communities. Environmental Microbiology 17: 4873–4884.

Merchant RE, Phillips TW, Udani J. 2015. Nutritional supplementation with *Chlorella pyrenoidosa* lowers serum methylmalonic acid in vegans and vegetarians with a suspected vitamin B12 deficiency. Journal of Medicinal Food 18: 1357–1362.

Neumann U, Derwenskus F, Gille A, Louis S, Schmid-Staiger U, Briviba K, Bischoff SC. 2018. Bioavailability and safety of nutrients from the microalgae *Chlorella vulgaris*, *Nannochloropsis oceanica* and *Phaeodactylum tricornutum* in C57BL/6 mice. Nutrients 10: 965.

Ohwada K, Taga N. 1973. Seasonal cycles of vitamin B12, thiamine and biotin in Lake Sagami. Patterns of their distribution and ecological significance. Internationale Revue der gesamten Hydrobiologie und Hydrographie 58: 851–871.

Ogle DH, Doll JC, Wheeler AP, Dinno A, 2025. FSA: Simple Fisheries Stock Assessment Methods. R package version 0.10.0

Panzeca C, Beck AJ, Tovar-Sanchez A, Segovia-Zavala J, Taylor GT, Gobler CJ, Sañudo-Wilhelmy SA. 2009. Distributions of dissolved vitamin B12 and Co in coastal and open-ocean environments. Estuarine, Coastal and Shelf Science 85: 223–230.

Parodi A, Leip A, De Boer IJM, Slegers PM, Ziegler F, Temme EHM, Herrero M, Tuomisto H, Valin H, Van Middelaar CE, et al. 2018. The potential of future foods for sustainable and healthy diets. Nature Sustainability 1: 782–789.

Pintner IJ, Altmeyer VL. 1979. Vitamin B12 binder and other algal inhibitors. Journal of Phycology 15: 391-398.

Provasoli L. 1958. Nutrition and ecology of protozoa and algae. Annual Review of Microbiology 12: 279–308.

Provasoli L, Carlucci AF. 1974. Vitamins and growth regulators. In Algal Physiology and Biochemistry (WDP Stewart, Ed.), pp. 741–787, Oxford: Blackwell Scientific Publications.

Raux E, Lanois A, Levillayer F, Warren MJ, Brody E, Rambach A, Thermes C. 1996. *Salmonella typhimurium* cobalamin (vitamin B12) biosynthetic genes: functional studies in *S. typhimurium* and *Escherichia coli*. Journal of Bacteriology 178: 753–767.

Ritz C, Baty F, Streibig JC, Gerhard D. 2015. Dose-response analysis using R. PLoS ONE 10: e0146021.

Sahni MK, Spanos S, Wahrman MZ, Sharma GM. 2001. Marine corrinoid-binding proteins for the direct determination of vitamin B12 by radioassay. Analytical Biochemistry 289: 68– 76.

Sañudo-Wilhelmy SA, Cutter LS, Durazo R, Smail EA, Gómez-Consarnau L, Webb EA, Prokopenko MG, Berelson WM, Karl DM. 2012. Multiple B-vitamin depletion in large areas of the. Proceedings of the National Academy of Sciences USA 109: 14041–14045.

Sañudo-Wilhelmy SA, Gómez-Consarnau L, Suffridge C, Webb EA. 2014. The role of B vitamins in marine biogeochemistry. Annual Review of Marine Science 6: 339–367.

Sayer AP, Llavero-Pasquina M, Geisler K, Holzer A, Bunbury F, Mendoza-Ochoa GI, Lawrence AD, Warren MJ, Mehrshahi P, Smith AG. 2024. Conserved cobalamin acquisition protein 1 is essential for vitamin B12 uptake in both *Chlamydomonas* and *Phaeodactylum*. Plant Physiology 194: 698–714.

Schauer K, Rodionov DA, de Reuse H. 2008. New substrates for TonB-dependent transport: do we only see the ‘tip of the iceberg’? Trends in Biochemical Sciences 33: 330–338.

Shelton AN, Seth EC, Mok KC, Han AW, Jackson SN, Haft DR, Taga ME. 2018. Uneven distribution of cobamide biosynthesis and dependence in bacteria predicted by comparative genomics. The ISME Journal 13: 789–804.

Sokolovskaya OM, Plessl T, Bailey H, Mackinnon S, Baumgartner MR, Yue WW, Froese DS, Taga ME. 2021. Naturally occurring cobalamin (B12) analogs can function as cofactors for human methylmalonyl-CoA mutase. Biochimie 183: 35-43.

Sokolovskaya OM, Shelton AN, Taga ME. 2020. Sharing vitamins: Cobamides unveil microbial interactions. Science 369: eaba0165.

Soto MA, Desai D, Bannon C, LaRoche J, Bertrand EM. 2023. Cobalamin producers and prokaryotic consumers in the Northwest Atlantic. Environmental Microbiology 25: 1300– 1313.

Stupperich E, Nexø E. 1991. Effect of the cobalt-N coordination on the cobamide recognition by the human vitamin B12-binding proteins intrinsic factor, transcobalamin and haptocorrin. European Journal of Biochemistry 199: 299–303.

Tang YZ, Koch F, Gobler CJ. 2010. Most harmful algal bloom species are vitamin B1 and B12 auxotrophs. Proceedings of the National Academy of Sciences USA 107: 20756–20761.

Tokuşoglu Ö, Ünal MK. 2003. Biomass nutrient profiles of three microalgae: Spirulina platensis, *Chlorella vulgaris* and *Isochrisis galbana*. Journal of Food Science 68: 1144–1148.

Vancaester E, Depuydt T, Osuna-Cruz CM, Vandepoele K. 2020. Comprehensive and functional analysis of horizontal gene transfer events in diatoms. Molecular Biology and Evolution 37: 3243–3257.

Watanabe F, Takenaka S, Kittaka-Katsura H, Ebara S, Miyamoto E. 2002. Characterization and bioavailability of vitamin B12-compounds from edible algae. Journal of Nutritional Science and Vitaminology 48: 325–331.

Wells ML, Potin P, Craigie JS, Raven JA, Merchant SS, Helliwell KE, Smith AG, Camire ME, Brawley SH. 2017. Algae as nutritional and functional food sources: revisiting our understanding. Journal of Applied Phycology 29: 949–982.

Wickham H, Averick M, Bryan J, Chang W, McGowan LD, François R, Grolemund G, Hayes A, Henry L, Hester J, Kuhn M, Pedersen TL, Miller E, Bache SM, Müller K, Ooms J, Robinson D, Seidel DP, Spinu V, Takahashi K, Vaughan D, Wilke C, Woo K, Yutani H. 2019. Welcome to the tidyverse. Journal of Open Source Software 4:1686.

Wienhausen G, Dlugosch L, Jarling R, Wilkes H, Giebel H-A, Simon M. 2022. Availability of vitamin B12 and its lower ligand intermediate α-ribazole impact prokaryotic and protist communities in oceanic systems. The ISME Journal 16: 2002–2014.

Wienhausen G, Noriega-Ortega BE, Niggemann J, Dittmar T, Simon M. 2017. The exometabolome of two model strains of the Roseobacter group: A marketplace of microbial metabolites. Frontiers in Microbiology 8: 1985.

Yi S, Seth EC, Men YJ, Stabler SP, Allen RH, Alvarez-Cohen L, Taga ME. 2012. Versatility in corrinoid salvaging and remodeling pathways supports corrinoid-dependent metabolism in *Dehalococcoides mccartyi*. Applied and Environmental Microbiology 78: 7745–7752.

