## Supplemental information for "Cellular basis of B_12_ uptake and remodelling in microalgae revealed using a novel bioassay"

### *New Phytologist* Supporting Information

Article acceptance date: Click here to enter a date.

The following Supporting Information is available for this article:

**Table S1 Strains used in this work, and maintenance and experimental conditions.** CCAP refers to the UKRI NERC Culture Collection of Algae and Protozoa (CCAP), based in Oban, UK ([www.ccap.ac.uk](http://www.ccap.ac.uk)) and EPSAG The Culture Collection of Algae at the University.

| **Identifier code** | **Species** | **Background** | **Reference** | **Growth media** | **Conditions** |
| --- | --- | --- | --- | --- | --- |
| WT-12 | *Chlamydomonas reinhardtii* | *Chlamydomonas reinhardtii* wild-type strain  12 (WT-12) nit^-^ (derived from strain CC124 137c mt^-^ nit1 nit2) | Davey *et al.* 2014 | Tris-acetate phosphate (TAP) medium with Kropat’s trace elements (Kropat *et al*., 2011) excluding selenium | 60-80 μmol photons m^-2^s^-1^ 16:8 light dark cycles at 110 rpm 25 °C |
| UVM4 | *Chlamydomonas reinhardtii* | NA | Neupert *et al.* 2009 | TAP medium with Kropat’s trace elements (Kropat *et al*., 2011) excluding selenium | 60-80 μmol photons m^-2^s^-1^ 16:8 light dark cycles at 110 rpm 25 °C |
| metE7 | *Chlamydomonas reinhardtii* | WT12 (derived from strain CC124 137c mt^-^ nit1 nit2) | Helliwell *et al*. 2015 | TAP medium with Kropat’s trace elements (Kropat *et al*., 2011) excluding selenium, with 200 ng/L cyanocobalamin | 60-80 μmol photons m^-2^s^-1^ 16:8 light dark cycles at 110 rpm 25 °C |
| metE4 | *Chlamydomonas reinhardtii* | UVM4 | Bunbury *et al*. 2020 | TAP medium with Kropat’s trace elements (Kropat *et al*., 2011) excluding selenium, with 200 ng/L cyanocobalamin | 60-80 μmol photons m^-2^s^-1^ 16:8 light dark cycles at 110 rpm 25 °C |
| IM4 | *Chlamydomonas reinhardtii* | UVM4 | Sayer *et al.* 2024 | TAP medium with Kropat’s trace elements (Kropat *et al*., 2011) excluding selenium | 60-80 μmol photons m^-2^s^-1^ 16:8 light dark cycles at 110 rpm 25 °C |
| IM4 comp | *Chlamydomonas reinhardtii* | IM4 | Sayer *et al.* 2024 | TAP medium with Kropat’s trace elements (Kropat *et al*., 2011) excluding selenium, with 200 ng/L cyanocobalamin | 60-80 μmol photons m^-2^s^-1^ 16:8 light dark cycles at 110 rpm 25 °C |
| metE4_cobTD5 KO | *Chlamydomonas reinhardtii* | metE4 | This work | TAP medium with Kropat’s trace elements (Kropat *et al*., 2011) excluding selenium | 60-80 μmol photons m^-2^s^-1^ 16:8 light dark cycles at 110 rpm 25 °C |
| Wild type (WT) CCAP1055/1 | *Phaeodactylum tricornutum* | NA | CCAP | F/2 media (Guillard *et al*. 1975) minus Silica & vitamins | 30-50 μmol photons m^-2^s^-1^ 16:8 light dark cycles at 110 rpm at 18 °C |
| ΔMETE-C9 | *Phaeodactylum tricornutum* | WT(1055/1) | This work | F/2 media (Guillard et al. 1975) minus Silica & thiamine, with 200 ng/L cyanocobalamin | 30-50 μmol photons m^-2^s^-1^ 16:8 light dark cycles at 110 rpm at 18 °C |
| ∆CBA1-3 | *Phaeodactylum tricornutum* | WT(1055/1) | Sayer *et al.* 2024 | F/2 media (Guillard *et al*. 1975) minus Silica & vitamins | 30-50 μmol photons m^-2^s^-1^ 16:8 light dark cycles at 110 rpm at 18 °C |
| WT CCAP 211/11B | *Chlorella vulgaris* | NA | CCAP | Modified Bold’s basal Medium (3N-BBM, Bischoff & Bold 1963) or TAP medium with Kropat’s trace elements (Kropat *et al*., 2011) excluding selenium | 60-80 μmol photons m^-2^s^-1^ 16:8 light dark cycles at 110 rpm 25 °C |
| WT (SAG 45/2) | *Lobomonas rostrata* | NA | EPSAG | TAP medium with Kropat’s trace elements (Kropat *et al*., 2011) excluding selenium, with 200 ng/L cyanocobalamin | 60-80 μmol photons m^-2^s^-1^ 16:8 light dark cycles at 110 rpm 25 °C |

**Table S2 Details of the cell density and concentration of B_12_ used for the uptake experiments.**

| **Species** | **Media** | **Concentration of B_12_ (pM)** | **Cell density (Cell/ml)** |
| --- | --- | --- | --- |
| *C. reinhardtii* | TAP | 150 | 5x10^6^ |
| *P. tricornutum* | F/2 minus vits+ silica | 600 | 5x10^6^ |
| *C. vulgaris* | 3N-BBM | 150 | 2.5x10^6^ |

**Table S3** **Details of oligonucleotides used in this study.**

| **Name** | **Sequence (5'->3')** | **Info** |
| --- | --- | --- |
| gMETEin.fwd | CACAGTGGGTGATATGTACCTCTACG | Red Fig. S1 |
| gMETEin.rv | CATTTTTTCATGCCTATTCACATTAGC | Red Fig. S1 |
| NAT.fwd | GAGGTCACCAACGTCAACG | Blue Fig. S1 |
| NAT.rv | AGTGAACACGACGCTGAAGG | Blue Fig. S1 |
| gMETE.fwd | TATCCGCGAACGACCTACAG | Black Fig. S1 |
| gMETE.rv | GAACAGCGCGACTTTTTGG | Black Fig. S1 |
| ts113_COBTextr_fwd | TGCGGCCACCTGTCGACAAC |  |
| ts115_COBTextr_rev | AGTGGTGACAGCGCTGAACAC |  |
| ts101_sgRNA_guide_rev | AAAAGCACCGACTCGGTGCCACTTTTTCAAGTTGATAACGGACTAGCCTTATTTTAACTTGCTATTTCTAGCTCTAAAAC | Reverse primer to generate dsDNA template for *in-vitro* transcription |
| ts126_COBT_sguide8_fwd | GAAATTAATACGACTCACTATAGAAAAGCGGCGATGGACGCCAGTTTTAGAGCTAGAAATAGCAAG | Forward primer to generate dsDNA template for *in-vitro* transcription |
| ts127_COBT_Cas9_ssODN | GACGGCTGGCGCAGCGCATGAATGGGCGGCGGCGTACGCCCAGGCCTAGATAACTAGAATTCCCAAGGCGAAGCCTGTGGGCTCCCTAGGTGCGTACGTCAGTGTTCAGCGCT | ssODN homology template for KO |
| CobT Cas9 target site | AAAAGCGGCGAUGGACGCCAGUUUUAGAGCUAGAAAUAGCAAGUUAAAAUAAGGCUAGUCCGUUAUCAACUUGAAAAAGUGGCACCGAGUCGGUGC | spacer underlined |

**Table S4** **The parameters used to relate the growth of bioassay strain to the concentration of B12 using the DRC package in R (Ritz et al. 2015).**

| **Equation** | **Parameter** |
| --- | --- |
| B | Slope |
| D | Lower limit |
| A | Upper limit |
| C | ED50 |
| y | Optical density of sample |
| Equation $x=C \times{(\frac{(A-y)}{(y-D)})}^{(\frac{1}{B})}$ | |

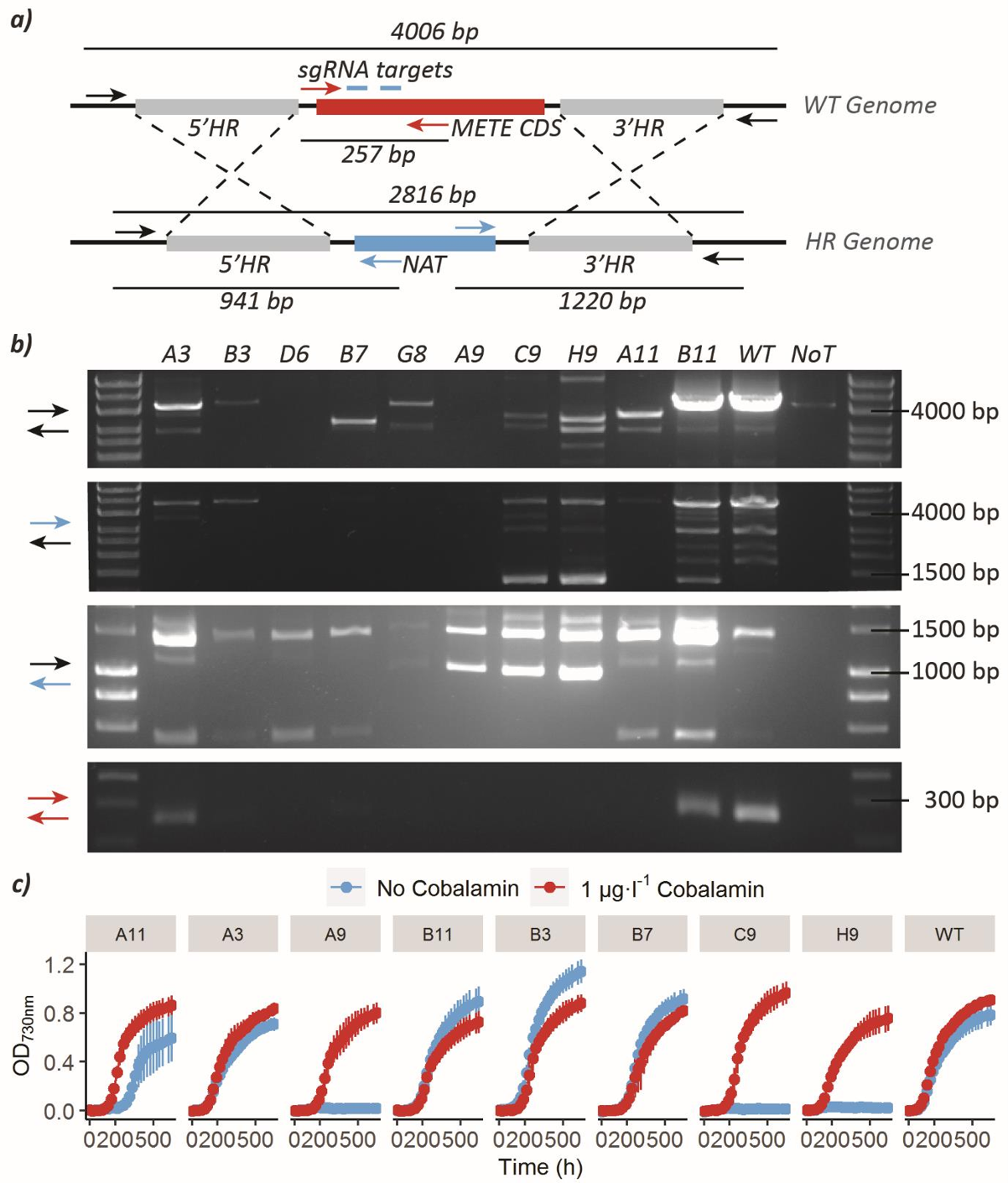

**Fig. S1 Genotyping and characterisation of CRISPR/Cas9 PtMETE mutants. a)** Depicts the genotyping strategy with 4 primer pairs (colour coded, Table S3) with expected band sizes indicated (associated with thin lines). **b)** PCR results for the four primer pairs in four different panels (colour coded on the left). The PCR for most primer pairs show many unspecific bands, probably explained by the repetitive nature of the METE gene. Colonies C9 and H9 show bands consistent with a biallelic homologous recombination knock-out genotype. **c)** Phenotypic characterization of 8 independent secondary colonies grown in triplicates with (1 μg·l^-1^) or without cobalamin supplementation in technical triplicate in 96- well plates and growth tracked as OD730nm for 20 days with a plate reader. Colonies A9, C9 and H9 show cobalamin auxotrophy consistent with their genotyping results*.*

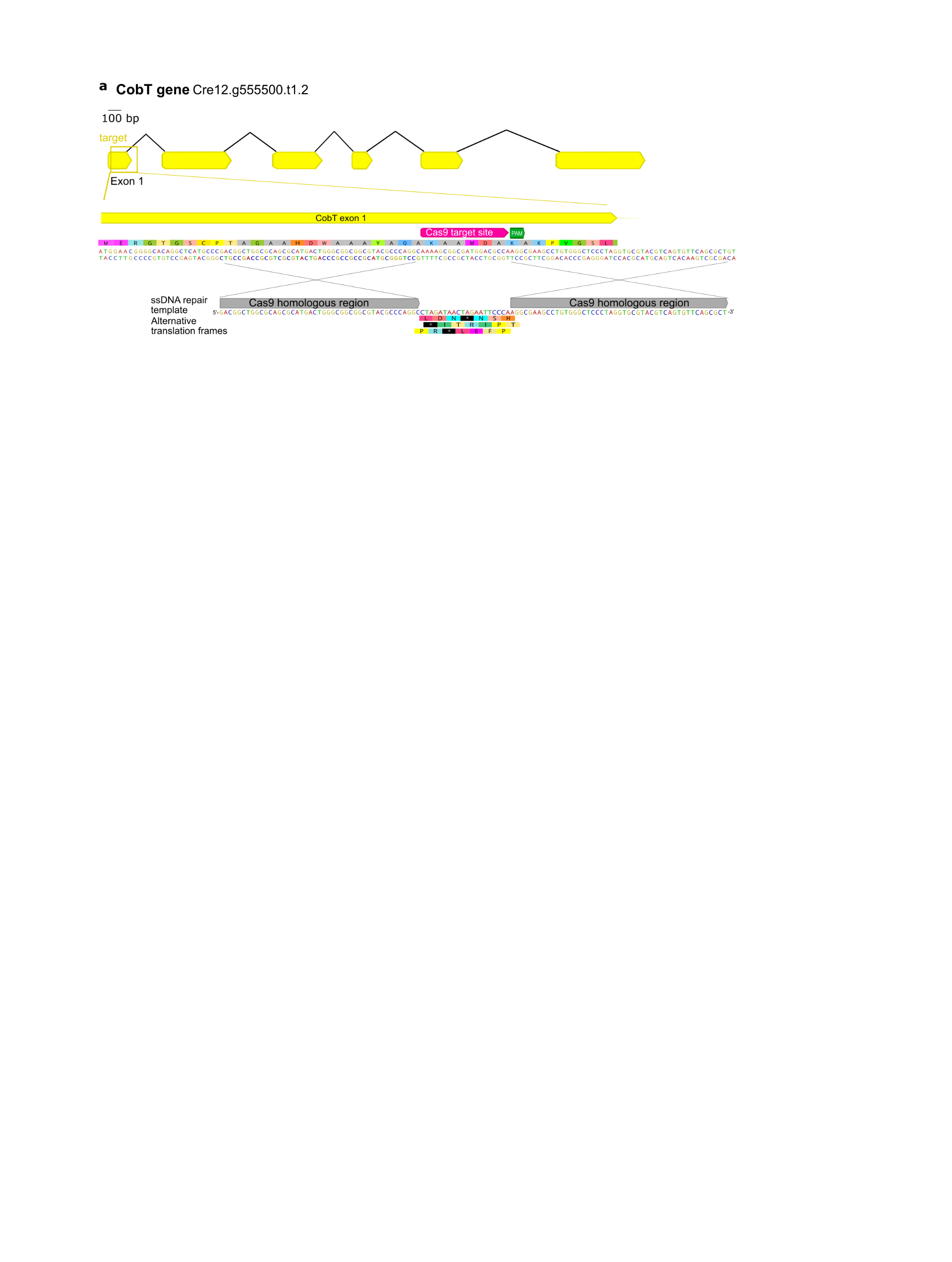

**Fig S2. Strategy for the generation of CrCOBT mutant.** Strategy to target COBT in metE4 background strain. Parts of this figure were generated with Geneious Prime 2022.1.1

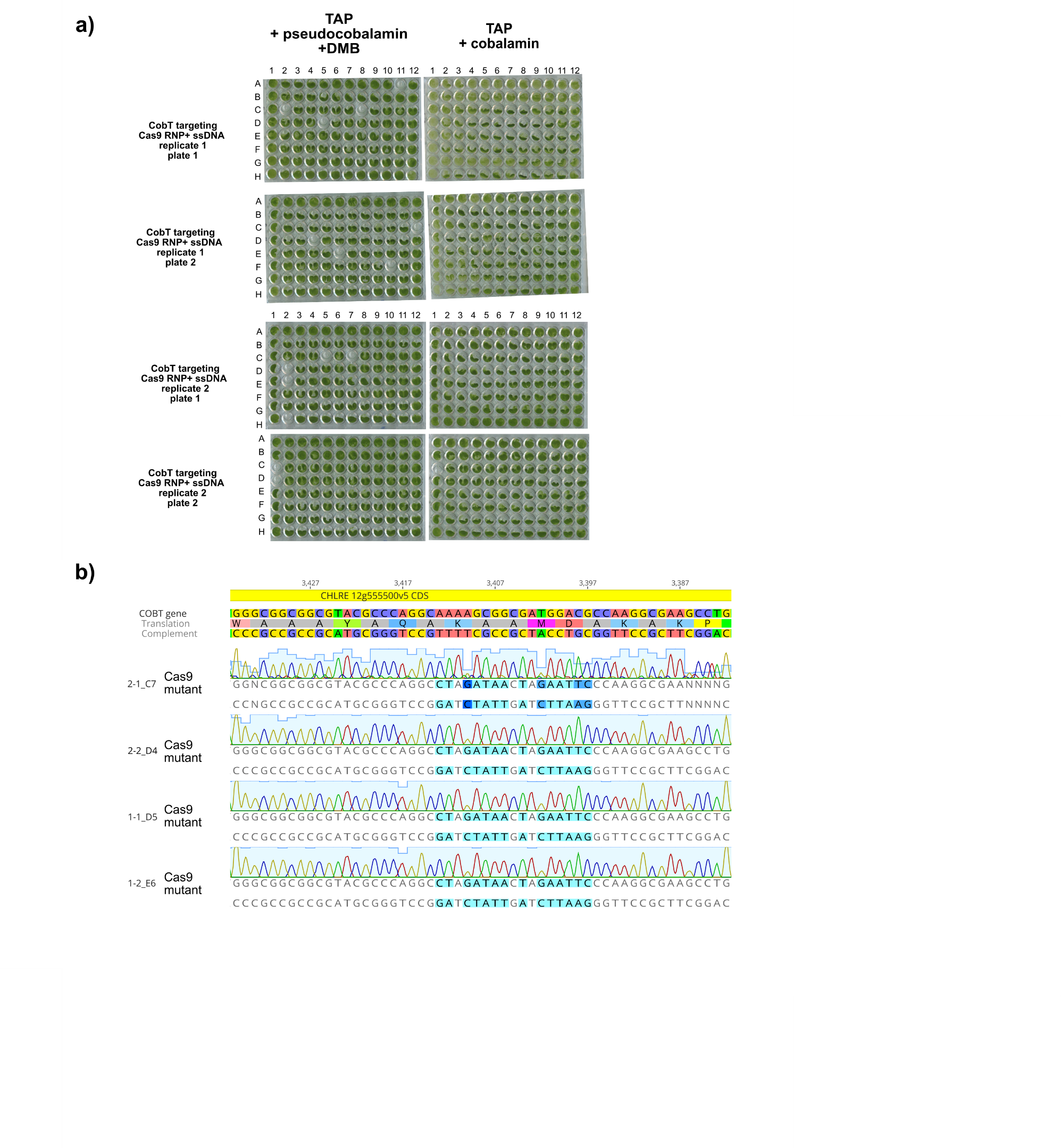
**Fig. S3 Identifying CrCOBT mutant. a)** shows mutant screening using the lack of remodelling ability, 4 mutants unable to grow when supplemented with pseudocobalamin +DMB were selected for genotyping, one from each plate shown. **b)** All 4 mutant strains show the intended edit, 1-1_D5 was taken forward and used in the experiment shown in Figure 5.

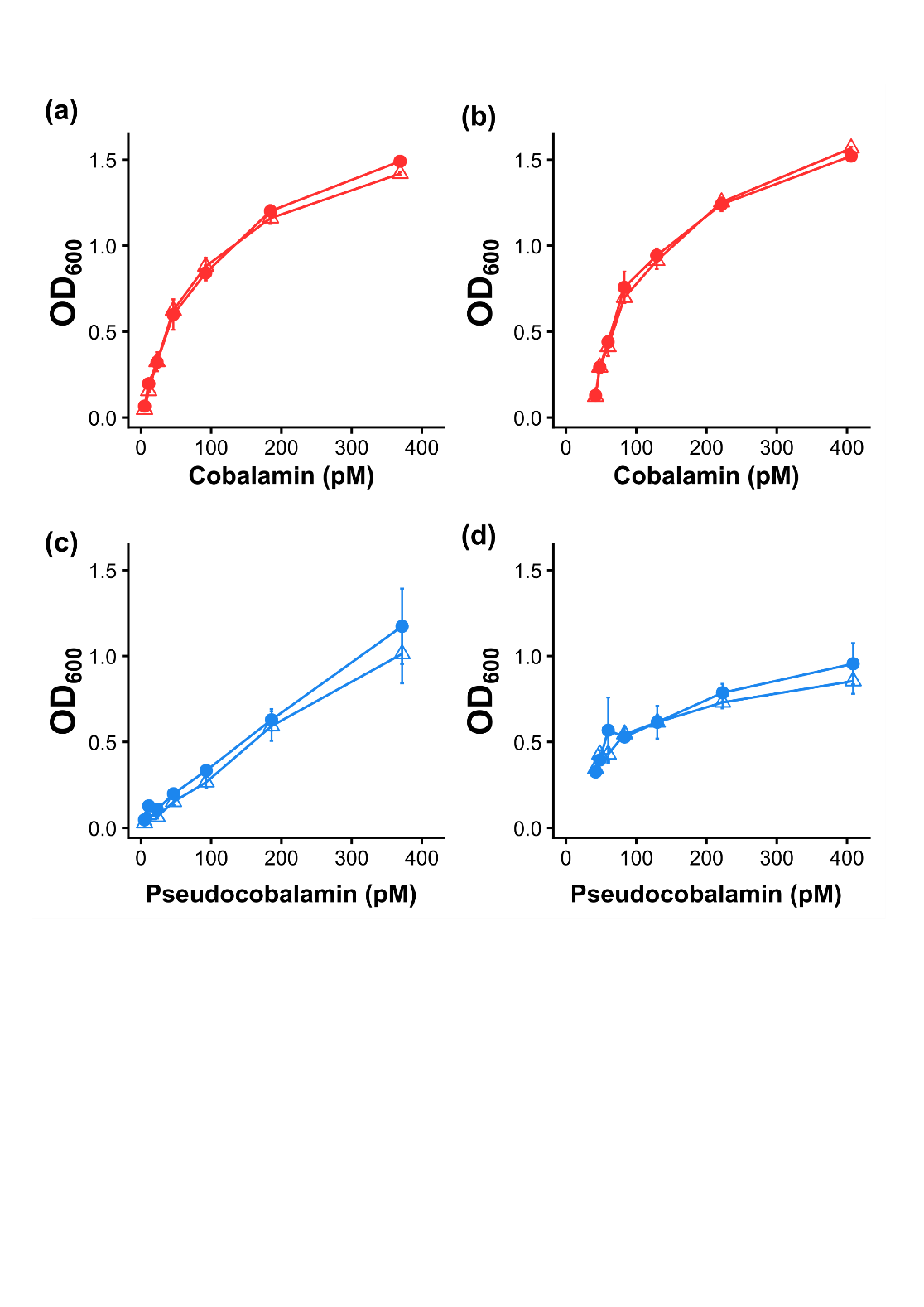

**Fig. S4 The growth of *Salmonella typhimurium* on mixtures of B_12_. a)** growth on various concentrations (pM) of cobalamin (Cbl, shown in red) with (triangles) or without (circles) 1 µM dimethyl-benzimidazole, DMB. **b)** the same concentrations of Cbl with the addition of 37 pM pseudocobalamin (PsCbl) with (triangles) or without (circles) DMB. **c)** shows the same experiment as (a) but with PsCbl (blue) only. **d)** PsCbl + 37 pM Cbl, with and without DMB. Optical density at 600 nm (OD600) used as a proxy for growth, after ~14-hour incubation at 37 °C. (n≥2, ±SD).

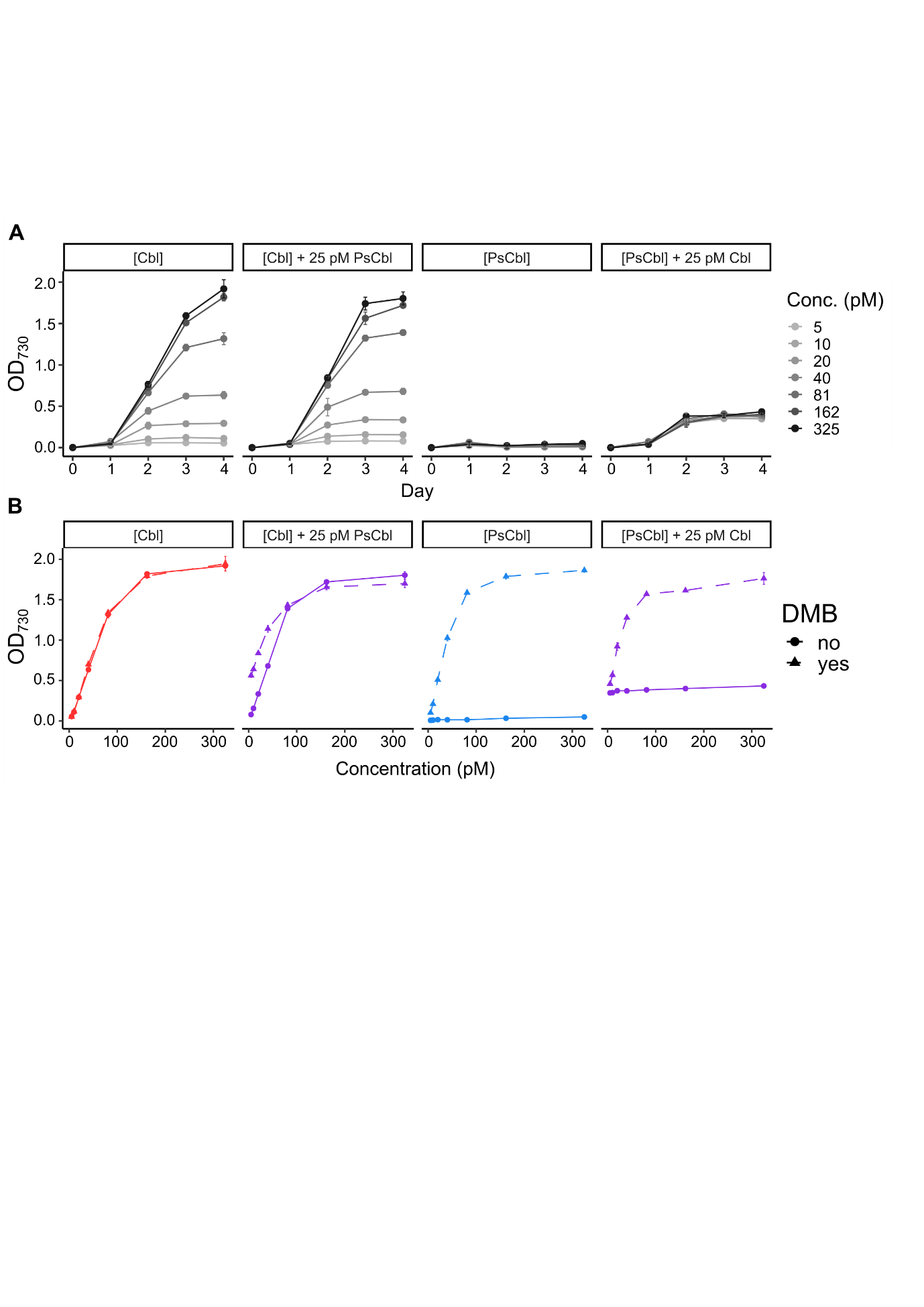

**Fig. S5 Effect of the presence of cobalamin and pseudocobalamin on the growth of *C. reinhardtii* metE7**. Growth on various concentrations (Conc. in pM) of B_12_ over 4 days, n=3 ±SD. Cbl = Cobalamin and PsCbl = pseudocobalamin.

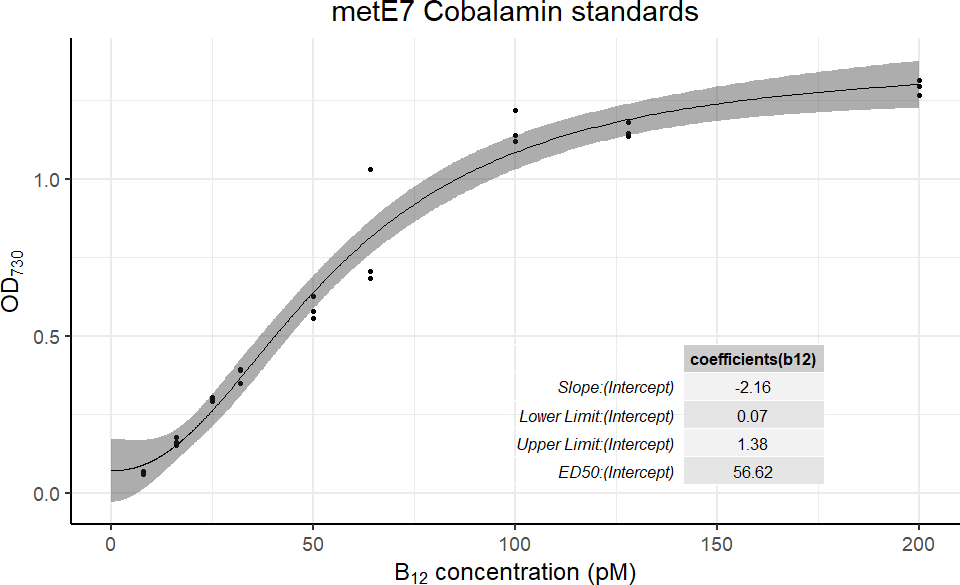

**Fig. S6 Example standard curve from *C. reinhardtii* metE7 bioassay.**

For each bioassay, separate standard curves were generated using the same culture for inoculum to mitigate any variability between starting cell physiology. The figures below show B_12_ concentrations calculated using curves generated from data in Fig. 3 and the logistic equations. Average of 3 replicates.

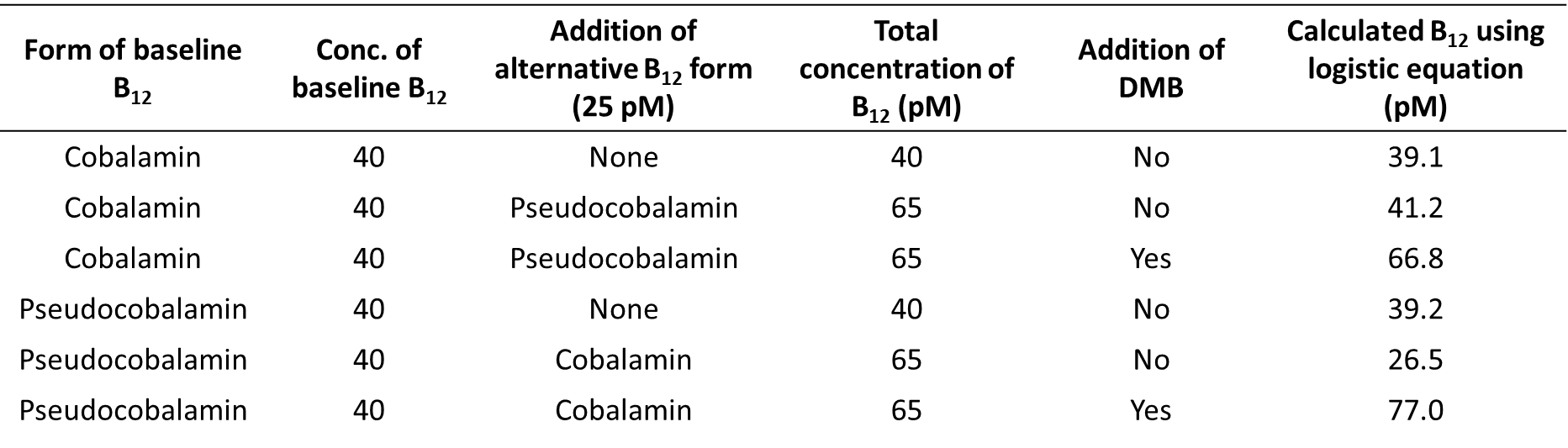

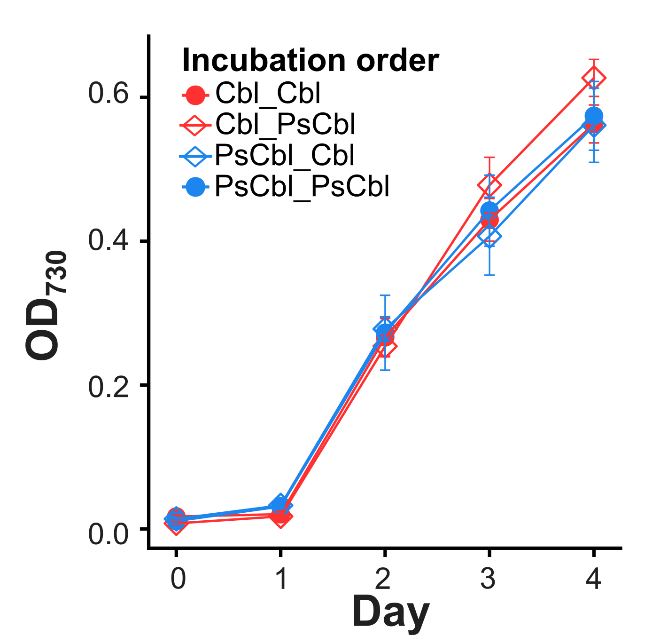

**Fig. S7 Confirmation of *C. reinhardtii* metE7 cell viability after sequential incubation with B_12_ analogues**. 200 pM cobalamin was resupplied to all replicates after sequential incubation with different combinations of cobalamin (Cbl) and pseudocobalamin (PsCbl), optical density at 730 nm used as a proxy for growth (OD_730_) cultures grown for 4 days (n=4, mean ±SE).

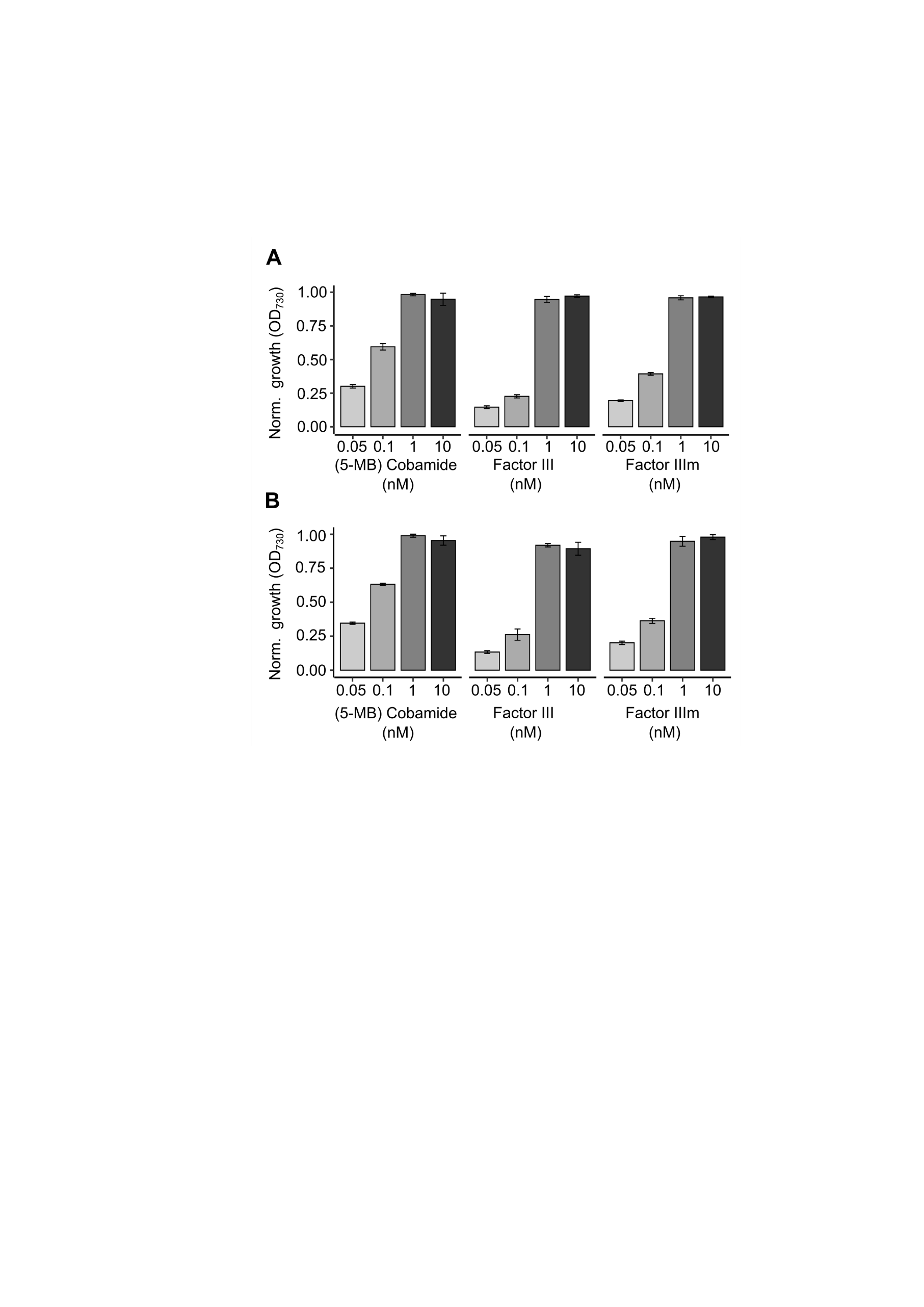
**Fig. S8 *C. reinhardtii* metE4 (A) and COBT knockout line (B) growth on 3 other benzimidazole B_12_ analogues.** Both strains were grown in liquid media for 4-days. Growth measured using optical density at 730 nm and normalised by dividing all values by the highest optical density achieved by each strain. (n=3, mean ±SD).

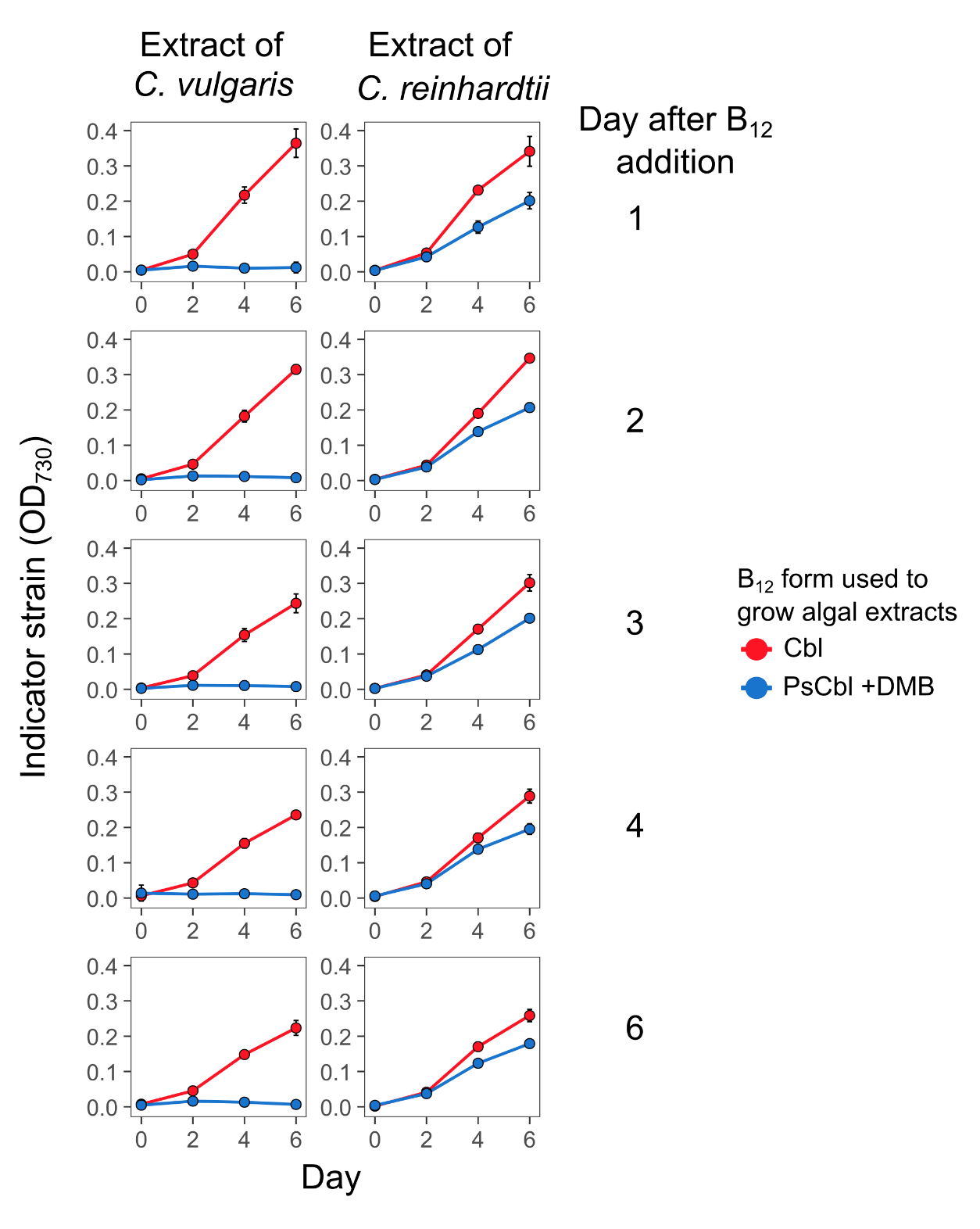

**Fig. S9 The growth of indicator strain *L. rostrata* on extracts of *C. vulgaris* or *C. reinhardtii* which in turn were grown with cobalamin or pseudocobalamin + DMB.** Cultures of *C. vulgaris* strains and *C. reinhardtii* were inoculated and grown for 2 days, then 250 pM B_12_ was added to each culture with 5 nM DMB added to the pseudocobalamin. To generate the extracts, 1 ml of culture was removed from each flask, and separated into cell and media fractions, cell fraction was boiled and added to medium used to grow *L. rostrata* cultures*.* Growth of *L. rostrata* monitored using optical density at 730 nm (OD730). n=3 ±SD. Day 4 shown in Figure 7.

**Methods S1**

**Equations necessary for the bioassay. Parameters explained in Table S4**

**Equation 1**

$$f\left( x \right)=C+(D+C)(1+exp(B\left( \log\left( x \right)-\log\left( e \right) \right))$$

**Equation 2**

$$x=C \times{(\frac{(A-y)}{(y-D)})}^{(\frac{1}{B})}$$

### References

Bischoff, H. W. & Bold, H. C. (1963), ‘Phycological Studies IV. Some soil algae from Enchanted Rock and related algal species’. U. Texas Pub. No. 6318. – adapted for CCAP <https://www.ccap.ac.uk/wp-content/uploads/MR_3N_BBM_V.pdf>

Bunbury, F., Helliwell, K. E., Mehrshahi, P., Davey, M. P., Salmon, D. L., Holzer, A., Smirnoff, N. and Smith, A. G. (2020) ‘Responses of a newly evolved auxotroph of *Chlamydomonas* to B_12_ deprivation’, *Plant Physiology*, 183(5), pp. 167–178. doi: 10.1104/pp.19.01375.

Davey, M.P., Horst, I., Duong, G.H., Tomsett, E.V., Litvinenko, A.C., Howe, C.J. and Smith, A.G., (2014), ‘Triacylglyceride production and autophagous responses in *Chlamydomonas reinhardtii* depend on resource allocation and carbon source.’ *Eukaryotic cell*, *13*(3), pp.392-400.

Helliwell, K. E., Collins, S., Kazamia, E., Purton, S., Wheeler, G. L. and Smith, A. G. (2015) ‘Fundamental shift in vitamin B_12_ eco-physiology of a model alga demonstrated by experimental evolution’, *The ISME Journal*, 9(6), pp. 1446–1455. doi: 10.1038/ismej.2014.230

Guillard, R. R. L. (1975) ‘Culture of Phytoplankton for Feeding Marine Invertebrates’, In Smith, W.L. and Chanley, M.H. (eds.) Culture of Marine Invertebrate Animals. Plenum Press New York, USA doi: 10.1007/978-1-4615-8714-9_3.

Kropat, J., *et al*. (2011) ‘A Revised Mineral Nutrient Supplement Increases Biomass and Growth Rate in *Chlamydomonas Reinhardtii*.’ *Plant J* 66(5): 770–80. <https://www.ncbi.nlm.nih.gov/pmc/articles/PMC3101321/pdf/nihms272702.pdf>

Neupert, J., Karcher, D., and Bock, R. (2009) ‘Generation of *Chlamydomonas* strains that efficiently express nuclear transgenes’, *Plant Journal*, 57(6), 1140–1150. doi: 10.1111/j.1365-313X.2008.03746.x.

Ritz, C., Baty, F., Streibig, J. C., Gerhard, D. (2015) ‘Dose-Response Analysis Using R’, *PLOS ONE*, 10(12), doi: 10.1371/journal.pone.0146021

Sayer, A.P., Llavero-Pasquina, M., Geisler, K., Holzer, A., Bunbury, F., Mendoza-Ochoa, G.I., Lawrence, A.D., Warren, M.J., Mehrshahi, P. and Smith, A.G., (2024) ‘Conserved cobalamin acquisition protein 1 is essential for vitamin B_12_ uptake in both *Chlamydomonas* and *Phaeodactylum*.’, *Plant Physiology*, *194*(2), pp.698-714.
